# Structure, dynamics and assembly of the ankyrin complex on human red blood cell membrane

**DOI:** 10.1101/2022.02.10.480008

**Authors:** Xian Xia, Shiheng Liu, Z. Hong Zhou

**Affiliations:** Department of Microbiology, Immunology, and Molecular Genetics, University of California, Los Angeles, CA 90095, USA; California NanoSystems Institute, University of California, Los Angeles, CA 90095, USA

## Abstract

The cytoskeleton of red blood cell (RBC) is anchored to cell membrane by the ankyrin complex. This complex is assembled during RBC genesis and comprises primarily band 3, protein 4.2 and ankyrin, whose mutations contribute to numerous human inherited diseases. High-resolution structures of the ankyrin complex have been long sought-after to understand its assembly and disease-causing mutations. Here, we analyzed native complexes on human RBC membrane by stepwise fractionation. Cryo-electron microscopy structures of nine band 3-associated complexes reveal that protein 4.2 stabilizes the cytoplasmic domain of band 3 dimer. In turn, the superhelix-shaped ankyrin binds to this protein 4.2 via ankyrin repeats (ARs) 6-13 and to another band 3 dimer via ARs 17-20, bridging two band 3 dimers in the ankyrin complex. Integration of these structures with both prior and our biochemical data supports a model of ankyrin complex assembly during erythropoiesis and identifies interactions essential for mechanical stability of RBC.

## Main

The human red blood cell (RBC, or the erythrocyte) is the most abundant cell in our blood and the principal gas exchanger between O_2_ and CO_2_ in our bodies. Devoid of nucleus, RBC has been engineered for a wide range of medical applications^1^. RBC exhibits unusual biconcave disc shape and remarkable membrane mechanical stability, both of which are essential for cycling through the vasculature for O_2_-CO_2_ exchange. These properties are endowed by the RBC cytoskeleton, which is bridged to the RBC membrane by junctional complex and ankyrin complex^2^. The ankyrin complex, which primarily contains band 3, protein 4.2, ankyrin and Rh subcomplex, connects the cytoskeleton to the membrane through the cytoplasmic domain of band 3 (refs.^3,4^). Defects in these cytoskeleton and cytoskeleton-associated proteins are associated with numerous human hereditary diseases, such as hereditary spherocytosis, South Asian ovalocytosis and hereditary stomatocytosis^5,6^.

Mass spectrometry and biochemical analyses of complexes from detergent-treated RBC membranes have shown that band 3, protein 4.2 and ankyrin interact with one another^7,8^, and that ankyrin repeats (ARs) 13-24 of ankyrin bind to the cytoplasmic domain of band 3 (refs.^9–11^). Protein 4.2 is a peripheral membrane protein with N terminal myristoylation^12^ and shares significant homology to transglutaminases but lacks transglutaminase activity^13,14^. Crystal structures are available separately for the membrane domain and cytoplasmic domain of band 3 (refs.^15,16^), as well as ARs 13 to 24 of ankyrin^17^. However, there are no structures available for any full-length proteins, let alone for any complexes containing them. Consequently, our understanding regarding the molecular interactions underlying the assembly and disease-causing mutations of the ankyrin complex remains extremely limited. In this study, we obtained native band 3, band 3-protein 4.2 complex and ankyrin complex centered on band 3, protein 4.2 and ankyrin (Fig. 1a) by stepwise fractionation of the erythrocyte membrane. A total of nine near-atomic resolution structures with various subunits of band 3, protein 4.2 and ankyrin were determined by cryo-EM, unraveling details of their interactions for the first time. These structures, combined with both prior and our biochemical data and knowledge about disease-causing mutations, supports a model of ankyrin complex assembly during erythropoiesis and reveal the importance of these interactions in linking the cytoskeleton to the membrane in RBC.

**Fig. 1:**
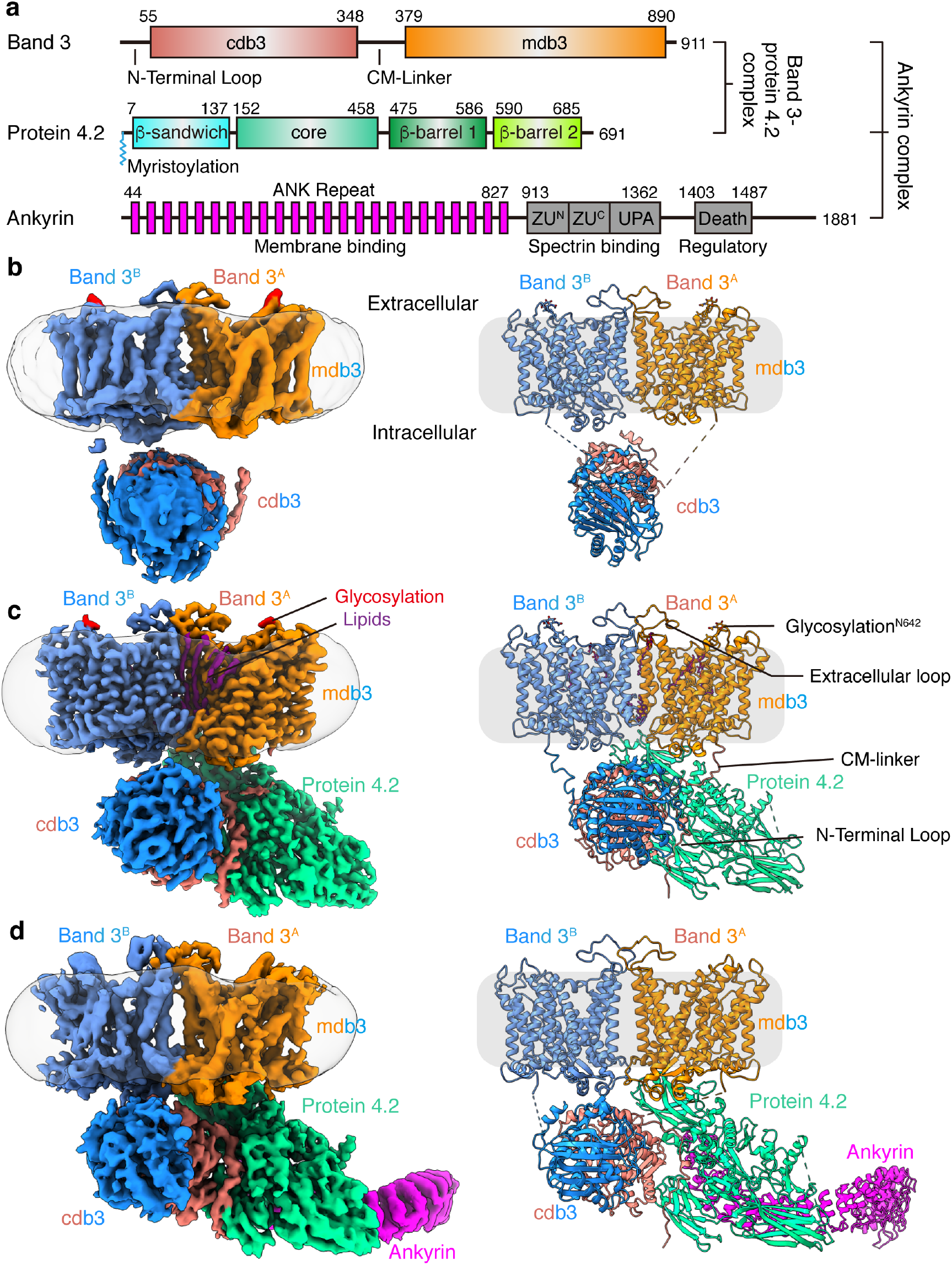
Cryo-EM structures of band 3, protein 4.2 complex and ankyrin-containing complex. (**a**) Schematic illustrating domain organizations of band 3, protein 4.2 and ankyrin. Residue numbers at domain boundaries are indicated. CM-linker in band 3 represents the linker between cdb3 and mdb3. The myristoylation site of protein 4.2 at the N-terminal residue Gly2 is indicated. (**b-d**) Cryo-EM maps and atomic models of the band 3 dimer (b), B_2_P_1_^diagonal^ complex (c) and B_2_P_1_A_1_ complex (d). The detergent belts are shown in transparent gray, depicting membrane boundaries. The maps in (c) and (d) are generated from focus refined maps combining membrane and cytoplasmic parts.

## Results

### Isolation of native band 3-associated complexes

To obtain the structures of full-length band 3 and related protein complexes, we analyzed the native proteins from human erythrocyte membrane by stepwise fractionation (Extended Data Fig. 1). Detergent solubilization of the erythrocyte membrane gave rise to three fractions: low-salt fraction, high-salt fraction 1, and high-salt fraction 2. To stabilize the ankyrin complex in the high-salt fraction 2 for cryo-EM, GraFix (Gradient Fixation)^18^ with glutaraldehyde was applied additionally.

SDS-PAGE and cryo-EM analyses both show that the predominant species in the low-salt fraction is band 3 (Extended Data Fig. 1b and 2c). The native band 3 is a dimer (Fig. 1b and Extended Data Fig. 2), and at an overall resolution of 4.8 Å, its structure reveals both the membrane domain (mdb3) and cytoplasmic domain (cdb3), each of which is similar to the crystal structures of mdb3 (ref.^16^) and cdb3 (ref.^15^), respectively. Though not resolved to high resolution, the cryo-EM density accommodates a cdb3 dimer structure with the characteristic reverse *V*-shape groove^19^, indicating that the structure of the band 3 dimer resolved here is in the *reversed-V* (*rev-V*) conformation (Extended Data Fig. 2g).

From the high-salt fraction 1, we identified 4 complexes—each containing a band 3 dimer (B_2_) and differently associated/oriented [either *loosely* associated, or *tightly* in *vertical* (75°) or *diagonal* (45°) orientation to the membrane] protein 4.2 as either monomer (P_1_) or dimer (P_2_)—designated as B_2_P_1_^loose^ (overall resolution 4.1 Å), B_2_P_1_^vertical^ (4.6 Å), B_2_P_2_^vertical^ (4.6 Å) and B_2_P_1_^diagonal^ (3.6 Å) (Table 1, Extended Data Fig. 3 and Movie S1). All of them have identical interaction between the cdb3 and protein 4.2. The majority (56%) of the complexes were the B_2_P_1_^diagonal^ complex (Fig. 1c and Extended Data Fig. 3c). Focused refinement further improved the resolution of the membrane (mdb3) and the cytoplasmic (cdb3 and protein 4.2) region to 3.3 Å and 3.1 Å, respectively (Extended Data Fig. 3c). Although loosely associated to band 3 based on its less robust density, protein 4.2 in B_2_P_1_^loose^ is oriented the same as it is in B_2_P_1_^vertical^. In the structure of B_2_P_2_^vertical^, the two protein 4.2 subunits do not interact with each other; rather, each subunit independently binds to a cdb3 of the band 3 dimer with two-fold symmetry.

**Table 1.** Cryo-EM data collection, refinement and validation statistics.

| **EMDB**<br>**(Focused refinement)** | **Band 3**<br>EMD-26148 | **B<sub>2</sub>P<sub>1</sub><sup>I</sup>**<br>EMD-26145 | **B<sub>2</sub>P<sub>1</sub><sup>V</sup>**<br>EMD-26146 | **B<sub>2</sub>P<sub>2</sub>**<br>EMD-26147 | **B<sub>2</sub>P<sub>1</sub><sup>d</sup>**<br>EMD-26142<br>(EMD-26143/<br>EMD-26144) | **B<sub>2</sub>P<sub>1</sub>A<sub>1</sub>**<br>EMD-26149<br>(EMD-26150) | **B<sub>2</sub>P<sub>1</sub>A<sub>2</sub>**<br>EMD-26151<br>(EMD-26152) | **B<sub>4</sub>P<sub>1</sub>A<sub>1</sub>**<br>EMD-26153 | **(B<sub>2</sub>P<sub>1</sub>A<sub>1</sub>)<sub>2</sub>**<br>EMD-26154 |
| --- | --- | --- | --- | --- | --- | --- | --- | --- | --- |
| **PDB** | 7TW2 |  | 7TW0 | 7TW1 | 7TVZ | 7TW3 | 7TW5 | 7TW6 |  |
| **Data collection and processing** |  |  |  |  |  |  |  |  |  |
| Magnification | 105K |  |  | 81K |  |  | 81K |  |  |
| Camera | K2 |  |  | K3 |  |  | K3 |  |  |
| Voltage (kV) | 300 |  |  | 300 |  |  | 300 |  |  |
| Electron exposure (e<sup>-</sup>/Å<sup>2</sup>) | 50 |  |  | 50 |  |  | 50 |  |  |
| Defocus range (μm) | -1.8 to -2.6 |  |  | -1.8 to -2.6 |  |  | -1.8 to -2.6 |  |  |
| Pixel size (Å) | 1.062 |  |  | 1.1 |  | 1.1 | 1.1 | 2.2 | 2.2 |
| Symmetry imposed | C1 | C1 | C1 | C2 | C1 | C1 | C1 | C1 | C1 |
| Particle number | 530K | 446K | 104K | 104K | 962K | 384K | 63K | 322K | 53K |
| Map resolution | 4.8 | 4.1 | 4.6 | 4.6 | 3.6<br>(3.3/3.1) | 4.4<br>(4.1) | 5.7<br>(4.4) | 5.6 | 8.5 |
| FSC threshold | 0.143 | 0.143 | 0.143 | 0.143 | 0.143 | 0.143 | 0.143 | 0.143 | 0.143 |
| **Refinement** |  |  |  |  |  |  |  |  |  |
| Map sharpening B-factor (Å<sup>2</sup>) | -237 |  | -192 | -212 | -190 | -214 | -324 |  |  |
| Model resolution (Å) | 4.1 |  | 3.9 | 3.8 | 3.5 | 4.4 | 5.6 | 5.5 |  |
| FSC threshold | 0.143 |  | 0.143 | 0.143 | 0.143 | 0.143 | 0.143 | 0.143 |  |
| Model composition |  |  |  |  |  |  |  |  |  |
| Non-hydrogen atoms | 12,838 |  | 18,266 | 23,625 | 18,659 | 22,641 | 27,048 | 27,184 |  |
| Protein residues | 1,619 |  | 2,309 | 2,993 | 2,340 | 2,944 | 3,579 | 3,515 |  |
| Ligand | 2 |  | 2 |  | 9 |  |  |  |  |
| B factors (Å<sup>2</sup>) |  |  |  |  |  |  |  |  |  |
| Protein | 457.5 |  | 196.0 | 191.9 | 68.9 | 276.3 | 258.2 | 592.2 |  |
| Ligand | 306.2 |  | 196.9 |  | 74.0 |  |  |  |  |
| R.m.s. deviations |  |  |  |  |  |  |  |  |  |
| Bond lengths (Å) | 0.003 |  | 0.003 | 0.003 | 0.002 | 0.004 | 0.003 | 0.004 |  |
| Bond angles (°) | 0.754 |  | 0.729 | 0.744 | 0.500 | 0.748 | 0.769 | 0.759 |  |
| Validation |  |  |  |  |  |  |  |  |  |
| MolProbity score | 1.56 |  | 1.33 | 1.47 | 1.34 | 1.68 | 1.84 | 1.68 |  |
| Clashscore | 9.46 |  | 5.6 | 7.46 | 4.08 | 9.8 | 12.0 | 10.2 |  |
| Poor rotamers (%) | 0.07 |  | 0 | 0 | 0 | 0.08 | 0 | 0.03 |  |
| Ramachandran plot |  |  |  |  |  |  |  |  |  |
| Favored (%) | 97.8 |  | 97.9 | 97.7 | 97.2 | 97.0 | 96.7 | 97.2 |  |
| Allowed (%) | 2.2 |  | 2.1 | 2.3 | 2.8 | 3.0 | 3.3 | 2.8 |  |
| Disallowed (%) | 0 |  | 0 | 0 | 0 | 0 | 0 | 0 |  |

Two ankyrin-bound complexes were obtained from the high-salt fraction 2, both containing B_2_P_1_^diagonal^ but with either one or two ankyrin molecules, which we designate as B_2_P_1_A_1_ (4.4 Å) or B_2_P_1_A_2_ (5.7 Å), respectively (Fig. 1d, Table 1 and Extended Data Fig. 4). Reprocessing the same particles extracted with a larger box size enabled visualization of ankyrin assembled on two dimers of band 3 (B_4_), which we designate as B_4_P_1_A_1_ and (B_2_P_1_A_1_)_2_ (Extended Data Fig. 4c).

In total, we isolated nine native band 3-associated complexes. Interactions among subunits of the band 3-associated complexes and related mutations, including those that cause human diseases, are detailed below.

### Structure of the full-length band 3

Previous efforts to obtain a full-length band 3 structure have not been fruitful. We found that, in the absence of protein 4.2, native full-length band 3 existed as a dimer in the low-salt fraction, and only the membrane domains were well resolved (Fig. 1b). This low-resolution nature of cdb3 is consistent with the previous observations that the CM-linker that bridge the cytoplasmic and the anchored membrane domains is flexible^19–21^. Among the nine band 3-associated complexes mentioned above, B_2_P_1_^diagonal^ has the highest resolution, resolving both mdb3 and cdb3 domains, as well as their linker, suggesting that binding of protein 4.2 restricts the relative movement of mdb3 and cdb3. Since B_2_P_1_^diagonal^ has the best resolution, subsequent description of band 3 structure will be based on this complex unless otherwise stated.

This first atomic structure of the native, full-length band 3 reveals long sought-after structural features. Compared to the domain crystal structures of mdb3 (ref.^16^) and cdb3 (ref.^15^), our native band 3 structure (Fig. 1c, Extended Data Fig. 5a and 6) is not only full-length, but also contains previously unresolved regions: N-terminal loop *(a.a.* 30-55), CM-linker (*a.a.* 350-369), residues 641-648 with the N-glycan site N642, a long external loop (*a.a.* 554-566), and the mdb3-bound lipids. The dimer interface of mdb3 is similar to that in the crystal structure. But in the dimer interface of the cryo-EM maps, we resolved densities of several lipids or detergent molecules, which may facilitate the dimerization of band 3. There are no obvious interactions between the cdb3 and mdb3. Band 3 is a Cl^-^/HCO_3_^-^ exchanger and belongs to the SLC4 family^22^. In all our cryo-EM structures, mdb3 adopts an outward-facing conformation as that of the mdb3 crystal structure locked by the inhibitor H2DIDS^16^ with a root mean square deviation (RMSD) of 0.931 Å for 443 of its 475 Cα atoms (Extended Data Fig. 5a,b). The N-terminal ends of the two transmembrane helices (TM3 and TM10) face each other and create a positive dipole, which may provide the binding site for substrate anions (Extended Data Fig. 5d,e). Remarkably, near residues R730 and E681 in the helical dipole, densities for four putative water molecules were observed, three of which possibly delineate the substrate binding site, as also predicted from the structure of substrate-bound SLC4 (NDCBE^23^), SLC23 (UraA^24^) and SLC26 (BicA^25^) family transporters (Extended Data Fig. 5d,e). Consistent with this assignment, mutation of the residue R748 or E699 in murine band 3 (equivalent to R730 and E681 in human band 3, respectively) resulted in loss of Cl^-^/HCO_3_^-^ exchange^26–28^, highlighting the importance of these residues in anion transport activity. Intriguingly, one molecule of n-Dodecyl-β-D-Maltoside (DDM) was identified at the interface between the gate and core domain of mdb3 (Extended Data Fig. 5c). Unlike H2DIDS, the DDM molecule does not block the substrate binding site. Instead, it may lock the relative rocking movement between the gate and core that is responsible for Cl^-^/HCO_3_^-^ translocation, leading to the observed outward-facing conformation.

### Protein 4.2 and its interactions with band 3

Efforts to determine the structure of protein 4.2 have hitherto been hindered by difficulties in obtaining purified protein 4.2 in soluble form. The atomic model of protein 4.2 built from our cryo-EM structure of the B_2_P_1_^diagonal^ complex contains 663 of its 691 residues. It has a triangular shape, with the body of the triangle formed by the core domain and the three vertices each formed by a domain with an immunoglobulin (Ig)-like fold^29^ (Fig. 2A and Extended Data Fig. 7a-e). These domains are sequential in sequence, and their folds and architecture are both similar to those of the transglutaminase family of enzymes in the closed conformation^13,30–32^ (RMSD of 3.898 Å with PDB 1L9N across 651 Cα atom pairs) (Fig. 2b). Therefore, we will use the transglutaminase domain names to describe corresponding domains of protein 4.2, *i.e.,* β-sandwich (*a.a.* 7-137, h-type Ig-like fold), core (*a.a.* 152-457), first (*a.a.* 475-586) and second (*a.a.* 590-685) β-barrel domains (s-type Ig-like fold), from N to C-terminus.

**Fig. 2:**
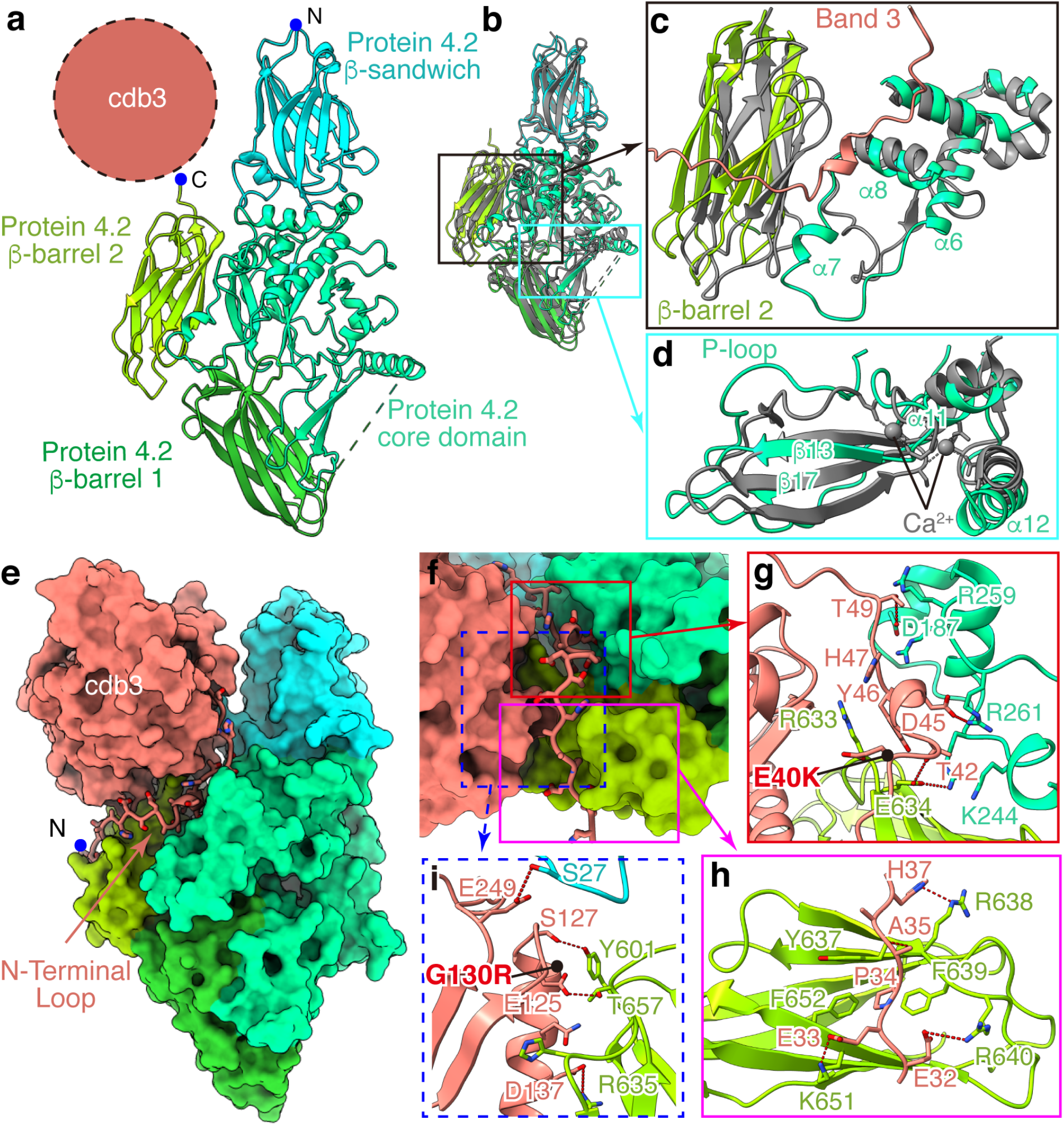
Protein 4.2 and its interactions with band 3. (**a**) Structure of protein 4.2 shown in ribbon. The position of cdb3 is indicated by the dashed circle. (**b**) Superposition of protein 4.2 with transglutaminase (gray, PDB: 1L9N) to identify structural differences in protein 4.2. (**c**) Movements of β-barrel 2 and α7 of core domain in protein 4.2. (**d**) Shifts of the P-loop and α12 in protein 4.2. (**e**) Interactions of the N-terminal loop of band 3 with protein 4.2. Protein 4.2 and cdb3 are shown as surface, while N-terminal loop of band 3 in ribbon and side chains are shown as sticks. (**f**) Enlarged view of the N-terminal loop in (e). Three important regions are boxed: region 1 (red box) indicates the interactions around residues 40 to 50 of N-terminal loop; region 2 (magenta box) indicates the interactions around residues 30 to 40 of N-terminal loop; region 3 (dashed blue box) shows the interactions between cdb3 and protein 4.2. (**g-i**) Details of the interactions in boxed regions of (f). Residues involved in the interactions are shown as sticks. Red dashed lines indicate hydrogen bonds and electrostatic interactions. Disease mutations (E40K and G130R) on band 3 are indicated by black dots.

The core domain has a globular shape with 10 helices (α3-12) flanking the 10 β strands (β10-19) that form three β-sheets in the middle (Extended Data Fig. 7a). Notably, the catalytic triads, C272, H330 and D353, in transglutaminase are replaced by A268, Q327 and H350, respectively (Extended Data Fig. 7b and 8); the three corresponding residues are more separated from each other in protein 4.2 than in transglutaminase (Extended Data Fig. 7b). The biochemically identified ATP-binding loop (P-loop, *a.a.* 316-322)^33^ in this domain is situated at its interface with β-barrel 1 (Fig. 2d), with a shift ~5 Å towards β-barrel 1 compared to the corresponding loop of transglutaminase (PDB: 1L9N)^31^. Intriguingly, a Ca^2+^ binding site in transglutaminase (PDB: 1L9N)^31^ corresponds to the N-terminus of helix α12 and the loop around β17 in the protein 4.2 core structure, both of which are closer to the β-barrel 1 domain than in transglutaminase (Fig. 2d). While no ATP and calcium were observed in our structure, these shifts suggest that ATP binding/hydrolysis or calcium binding may induce conformational changes to protein 4.2 (ref.^30^), possibly modulating interactions with ankyrin (see below).

The three Ig-like domains share typical β-strand topology, but the N-terminal proximal one (β-sandwich) contains one additional strand (d) and two short helices (Extended Data Fig. 7c-e). Upon binding of cdb3, β-barrel 2 and the region near α7 (*a.a.* 227-268) shift towards band 3 (Fig. 2c). The membrane-proximal region of protein 4.2 is connected to the density of detergent micelles and contains predominantly positively charged residues (Extended Data Fig. 7f). These structural observations are consistent with previous observations that protein 4.2 is myristoylated at the N-terminal Glycine residue and anchored to the lipid bilayer^12,34^.

Among the eight protein 4.2-containing structures, the interactions between band 3 and protein 4.2 are nearly identical. On band 3, this interaction is essentially through its cytoplasmic domain. The N-terminal loop (*a.a.* 30-55) was disordered in the absence of protein 4.2 and became ordered and visible upon binding protein 4.2 (Fig. 2e). The binding interface between band 3 and protein 4.2 can be divided into three regions (Fig. 2f). In region 1, a short helix and nearby loops of band 3 are docked into the groove formed by the core and the β-sandwich domains of protein 4.2 (Fig. 2g). The interactions include hydrogen bonds (band 3-protein 4.2: T42-E634, T49-D187) and electrostatic interaction between band 3 D45 and protein 4.2 R261, as well as hydrophobic interaction requiring band 3 Y46. In region 2, the band 3 loop (*a.a.* 30-39) lies on the surface of β-barrel 2 of protein 4.2. This interaction is mostly mediated by electrostatic interactions and hydrogen bonds (band 3-Protein 4.2: E32-R640, E33-K651, A35-R638, H37-S636), while the hydrophobic interaction involving P34 of band 3 also strengthens the interaction (Fig. 2h). In region 3, residues around α3 (*a.a.* 120-150) and the loop after β7 (*a.a.* 249-250) in cdb3 hydrogen-bond with protein 4.2’s β-barrel 2 domain and the β-sandwich domain, respectively (Fig. 2i). Besides the major interface described above, the CM-linker of band 3 interacts with the core domain of protein 4.2, further stabilizing the B_2_P_1_^diagonal^ complex (Extended Data Fig. 9f-h), which is not observed in the B_2_P_1_^vertical^ and B_2_P_2_^vertical^ complexes. Notably, protein 4.2 does not interact with mdb3 specifically; rather, the mdb3 proximal N terminus of protein 4.2 can restrict the rocking movement between the gate and core domains of mdb3 needed for ion exchange/transport. Such restriction might be the reason why band 3’s capability of ion transport decreases after binding protein 4.2 (ref.^35^). These close interactions between band 3 and protein 4.2 are consistent with both biochemical observations and disease-causing mutations. Protein 4.2 can be purified from the membrane only by relatively harsh treatments^36,37^. Two hereditary spherocytosis mutations (E40K and G130R)^38,39^ in cdb3, which lead to disproportionate loss of protein 4.2, are located on these binding interfaces (Fig. 2g,i), further highlighting the essential role of band 3-protein 4.2 interaction in the function of erythrocytes.

### Anchorage of ankyrin to protein 4.2 and band 3

Previous crystal structures of recombinantly-expressed ankyrin fragments indicated that the 89 kDa N-terminal membrane binding domain of erythrocyte ankyrin consists of 24 ankyrin repeats of approximately 33 amino acids each^17,40^. ARs 6-24 were modeled in our cryo-EM density maps of the four ankyrin-containing complexes [B_2_P_1_A_1_, B_2_P_1_A_2_, B_4_P_1_A_1_ and (B_2_P_1_A_1_)_2_] (Fig. 3a and Extended Data Fig. 10a,b,e). No significant global structural changes occur in either band 3 or protein 4.2 in these complexes upon ankyrin binding.

**Fig. 3:**
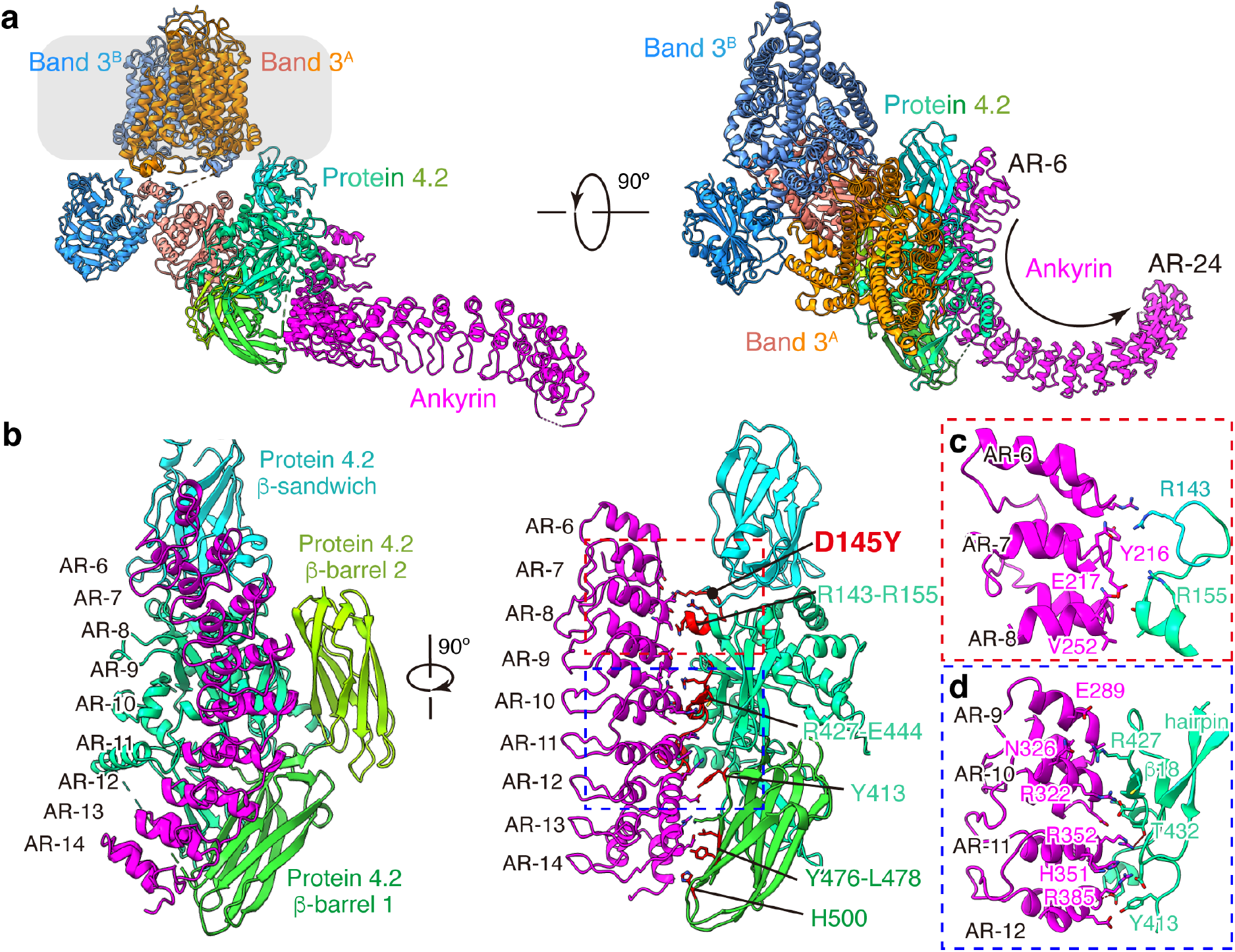
Anchorage of ankyrin to protein 4.2. (**a**) Atomic model of the B_2_P_1_A_1_ complex shown as ribbon. Approximate boundaries of the membrane are indicated in transparent gray. (**b**) Different views of the protein 4.2-ankyrin interface. Band 3 and ARs 15-24 of ankyrin are omitted for clarity. Residues of protein 4.2 involved in ankyrin interaction are colored in red and labeled. A disease mutation (D145Y) on protein 4.2 is indicated by a black dot. (**c-d**) Details of interactions in boxed regions of (b).

The interactions between ankyrin and protein 4.2 are the same in all four ankyrin-containing complexes; therefore, we focused our description below on B_2_P_1_A_1_, which has the best resolution (4.1 Å) for the region involving ankyrin-protein 4.2 interactions. Protein 4.2 interacts extensively with ARs 6-13 of ankyrin via a conserved surface on its core and β-barrel 1 domains (Fig. 3b, Extended Data Fig. 7h and 8). Overall, these two domains clamp ankyrin, with major binding sites located in the core domain of protein 4.2. Specifically, residues around α3 (*a.a.* R143-E152) of protein 4.2 interact with ARs 6-8 of ankyrin via hydrogen bonds, whereas residues between β17 and α12 (*a.a.* Y413-E439) contact extensively with ARs 9-12 through both hydrogen bonds and hydrophobic interactions (hydrophobic stacking between Y413 of protein 4.2 and R385, H351 of ankyrin) (Fig. 3c,d and Extended Data Fig. 8). This ankyrin-protein 4.2 binding is further strengthened by contacts between the β-barrel 1 of protein 4.2 (residues Y476-L478 and H500) and AR-13 of ankyrin (Fig. 3b). The previously identified hairpin of protein 4.2 (N163-D180) in the core domain^13,41^ has no interaction with band 3, but instead is close to AR-11 of ankyrin and thus may facilitate the association between protein 4.2 and ankyrin (Fig. 3d).

Besides interacting with protein 4.2, ankyrin also directly contacts band 3 in three [B_2_P_1_A_2_, B_4_P_1_A_1_ and (B_2_P_1_A_1_)_2_] of the four ankyrin-containing complexes (Fig. 4a and Extended Data Fig. 10a,b,e), specifically with ARs 17-20 binding cdb3. The details of this binding are illustrated with B_2_P_1_A_2_ (4.6 Å after focus classification) (Fig. 4b), including the following three sets of amino acids between ankyrin and band 3: AR-17 (L556-Q563) and AR-18 (*a.a.* R590-H596) with band 3 residues near α3 ( *a.a.* S127-G130) and β7 (*a.a.* E252-E257), AR-19 (*a.a.* R619-S627) with band 3 residues E151-K160 near a4, and AR-20 (*a.a.* K657-L663) with band 3’s loop residues K69-E72 and D183.

**Fig. 4:**
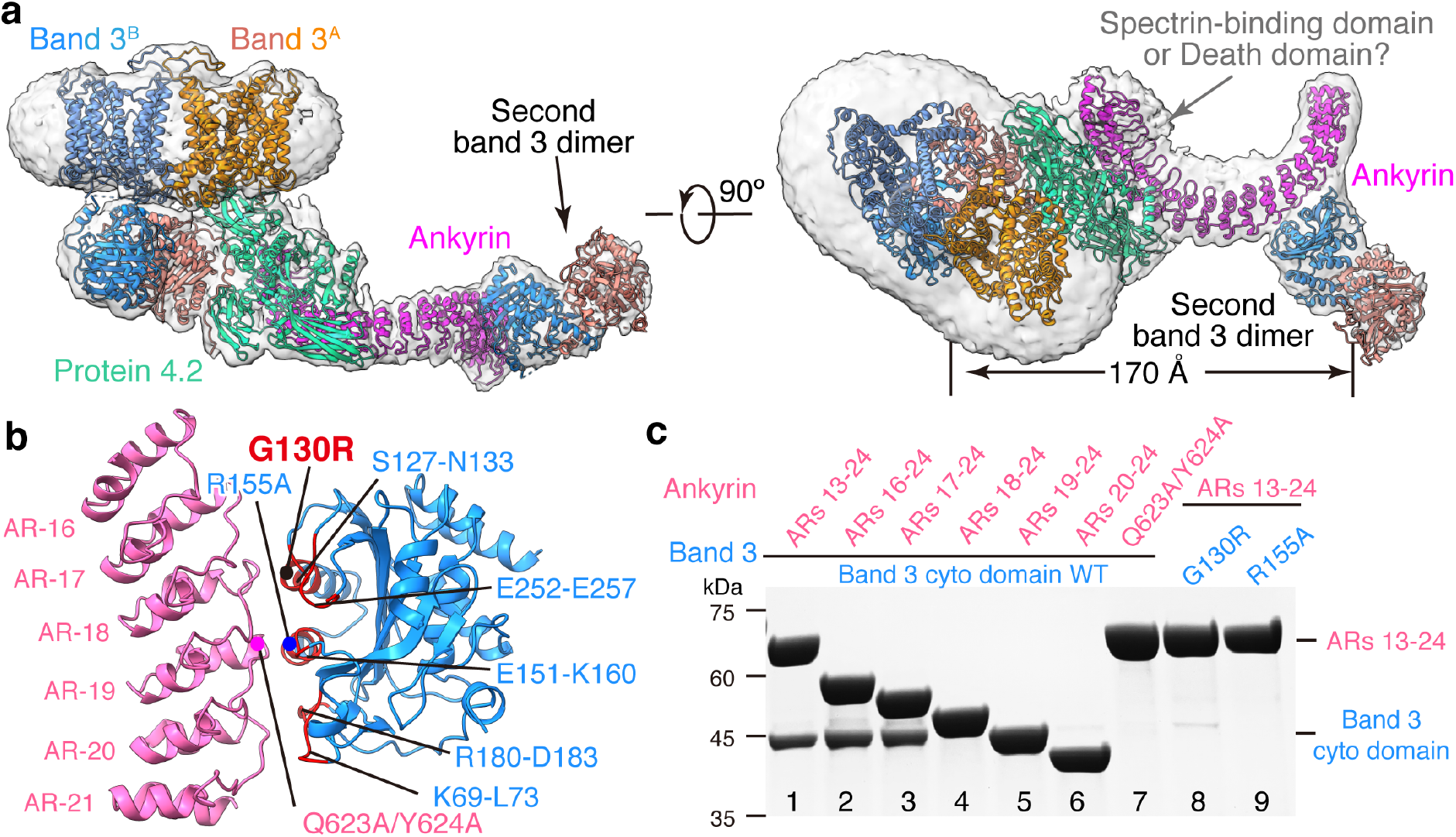
Interaction of ankyrin with band 3. (**a**) Two orthogonal views of the atomic model of B_4_P_1_A_1_ complex (ribbon), with density map in transparent gray. The density of the membrane domains of the second band 3 dimer are weak and only visible at low density threshold, and are thus not modeled. (**b**) Interaction between cdb3 and ankyrin from B_2_P_1_A_2_ complex. Residues of band 3 involved in ankyrin interaction are colored in red and labeled. Positions of the mutations used in (c) are indicated as dots. (**c**) SDS-PAGE gel of the His-tag pull-down assay from the recombinant proteins. Band 3 mutations G130R and R155A eliminated ankyrin binding; ankyrin truncations (ARs 18-24, ARs 19-24 and ARs 20-24) and the mutations on AR-19 (Q623A/Y624A) abolished band 3 interaction. The experiments were repeated independently for three times and representative results are shown here.

Biochemical data further confirmed these observed interactions between band 3 and ankyrin. First, our pull-down assay showed that the mutations G130R (disease mutation) on band 3 α3 and R155A on band 3 a4 eliminated ankyrin binding. In addition, ankyrin truncations ARs 18-24, ARs 19-24 and ARs 20-24, as well as mutations on AR-19 (Q623A/Y624A), abolished band 3 interaction (Fig. 4c), further validating the atomic details of the band 3-ankyrin interaction. Earlier, site-directed mutagenesis and antibody studies indicated that residues 63-73 (ref.^42^), 118-162 (ref.^9^) and 175-185 (ref.^42^) in the peripheral region of cdb3 were responsible for ankyrin binding. Site-directed spin-labeling on ankyrin demonstrated that its convex surface, other than its concave groove of ARs 18-20, served as primary sites to contact cdb3 (ref.^11^), which is consistent with our structures.

As indicated above, the protein 4.2 binding sites on band 3 partially overlap with the ankyrin binding sites on band 3 (Extended Data Fig. 6), indicating that protein 4.2 and ankyrin associate with band 3 monomer exclusively. The interface between protein 4.2 and ankyrin spans about 1630 Å^2^, which is ~2.4 fold larger than the interface of band 3-ankyrin (about 690 Å^2^). Combined with the described interactions between protein 4.2 and band 3 in the previous section, it can be inferred that protein 4.2 may function as a linker to strengthen the ankyrin-band 3 association, consistent with previous findings that protein 4.2 deficiency weakened ankyrin-band 3 association on the erythrocyte membrane^30,43,44^.

## Discussion

Prior biochemical analysis of band 3 multiprotein complex assembly during erythropoiesis established the temporal progression towards the assembly of the ankyrin complex from various sub-complexes, including band 3, band 3-protein 4.2 complex and Rh complex^44–46^. The ratio of abundance of band 3, protein 4.2, ankyrin in RBC is about 10:2:1 (ref.^47^), which differs from the stoichiometry ratio (4:1:1) of these proteins in the ankyrin complex. Therefore, as the most abundant protein on the mature RBC membrane^48^, band 3 could exist as a dimer without forming larger complexes with others in the mature RBC; and likewise, other sub-complexes could exist without forming the ultimate supra-complex with all components. Indeed, their existence on mature RBC is the basis for our ability to use the stepwise fractionation strategy to capture a total of nine native structures from the RBC membrane reported above though we could not rule out the possibility that some may have resulted from disassembly during isolation. These results now allow us to populate the previously depicted model of ankyrin complex assembly (Fig. 5; Movie S1) during erythropoiesis with experimentally observed sub-complex structures.

**Fig. 5:**
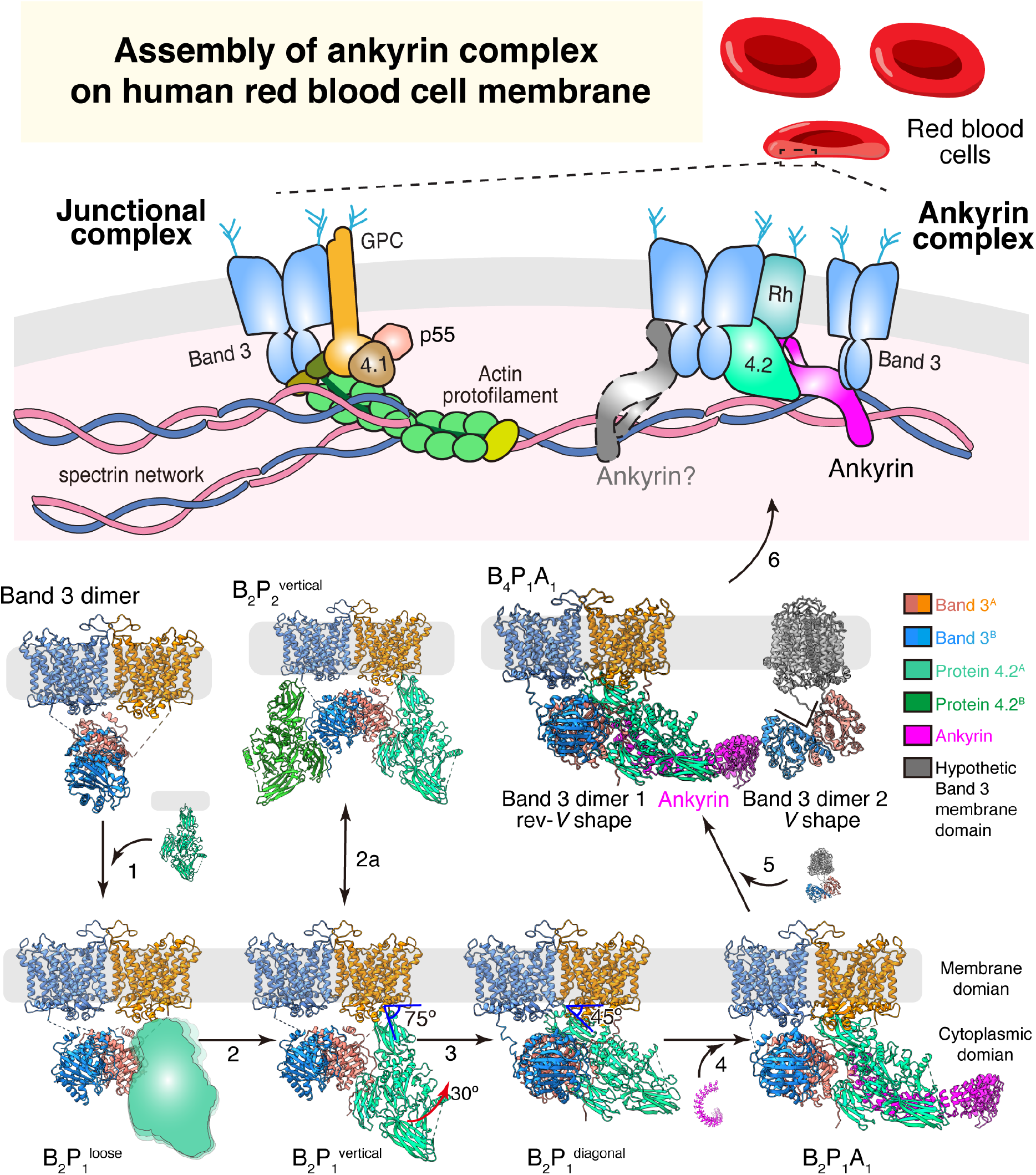
Schematic and possible assembly model of the ankyrin complex. Seven of the nine structures reported in the paper are depicted, each showing one possible assembly state (loosely-bound protein 4.2 and complexes involving the second ankyrin molecule are shown as cartoon). For the B_4_P_1_A_1_ complex, the membrane domains of the second band 3 dimer are hypothetically modelled (gray) to align to the membrane. Arrows depict possible directions of the assembly pathway, from the band 3 dimer to the ankyrin complex.

Assembly of ankyrin complex possiblely starts from free band 3 dimer^46^. By interacting with the *rev-V* shaped band 3, protein 4.2 is incorporated to form the band 3-protein 4.2 complex, first in a loosely-bound vertical conformation (B_2_P_1_^loose^, Fig. 5 step 1) and then converted into a tightly-bound vertical conformation (B_2_P_1_^vertical^) (Fig. 5 step 2 and Extended Data Fig. 3c). A second protein 4.2 molecule can further interact with the unoccupied cdb3, forming a C2-symmetric B_2_P_2_^vertical^ complex (Fig. 5 step 2a). Next, the membrane anchorage site of protein 4.2 moves from the edge to the center of the mdb3 dimer while cdb3 shifts off the 2-fold axis (Extended Data Fig. 9a-e), transitioning from B_2_P_1_^vertical^ into B_2_P_1_^diagonal^ (Fig. 5 step 3). Protein 4.2 interacts with the CM-linker of band 3 in B_2_P_1_^diagonal^, further stabilizing the diagonal conformation of protein 4.2 in the band 3-protein 4.2 complex (Extended Data Fig. 9f-h). Following the formation of B_2_P_1_^diagonal^, one ankyrin molecule binds to protein 4.2, resulting in the B_2_P_1_A_1_ complex (Fig. 5 step 4). By simultaneously interacting with protein 4.2 at ARs 6-13 and band 3 at ARs 17-20, ankyrin can bridge two band 3 dimers to form a B_4_P_1_A_1_ complex (Fig. 5 step 5), consistent with our results from gel-filtration analysis of the reconstituted ankyrin complex (Extended Data Fig. 10b) and the observation that binding of ankyrin to band 3 promoted the formation of band 3 tetramers^49,50^. To align mdb3 of both band 3 dimers to the cell membrane, the second band 3 dimer must be in the *V* shape conformation and without protein 4.2 binding (Fig. 5 step 5).

Notably absent from the above assembly picture are several other complexes, likely due to their flexibility and/or transient existence. For example, a second ankyrin molecule can bind to cdb3 not occupied by protein 4.2, forming a B_2_P_1_A_2_ complex (Extended Data Fig. 10c), which constitutes a small portion (14% of the particles) of ankyrin-bound complexes. Analytic gel-filtration analysis of the reconstituted ankyrin complex shows that the second ankyrin (Extended Data Fig. 10d) can be incorporated into the ankyrin complex. Furthermore, through the interaction between the ZU5-UPA domain of ankyrin and repeats 13-14 of β-spectrin^51^, the ankyrin complex links the spectrin network to the erythrocyte membrane (Fig. 5, step 6). Other complexes, such as the Rh complex, which contains RhCE, RhD, RhAG, CD47, LW and glycophorin B, can also interact with band 3, protein 4.2 and ankyrin^4,52^, forming the intact ankyrin complex. While association of protein 4.2 and ankyrin to band 3 may occur at early stage of erythropoiesis even prior to membrane integration, incorporation of the Rh complex is thought to happen afterwards on the cell membrane^45,46^. The validity of our proposed model of ankyrin complex assembly (Fig. 5) and other possible assembly intermediates during erythropoiesis await testing by cryo electron tomography of erythropoiesis at different stages.

The significance of the current study lies in both biology and technology perspectives. From the biology perspective, mutations on the components of the ankyrin complex can result in disorders in erythrocytes (hereditary spherocytosis, South Asian ovalocytosis, and hereditary stomatocytosis^5,6^) (Extended Data Fig. 10f). In hereditary spherocytosis, disruption of subunit interactions in the ankyrin complex results in loss of connection between the cytoskeleton and the membrane, consequently decreasing mechanical resistance and shortening the lifespan of the erythrocyte. Disease mutations, including G130R, E40K of band 3 and D145Y of protein 4.2, are located at the subunit binding interfaces. The availability of atomic structures of the ankyrin complex provides mechanic insight to red blood cell functions and paves the way for developing therapeutics against these diseases. From a technical perspective, the current work demonstrates an approach for direct visualization, at near-atomic resolution, of native protein complexes as they exist on membranes or in the cellular milieu. As such, notwithstanding obvious challenges in dealing with species only existing transiently in cells, this approach opens the door for structural study of native macromolecular complexes to capture their multiple conformational states (*e.g.,* changes in binding partners^53^ or cycling through sub-complexes like the spliceosome^54,55^) and during various functional stages (*e.g.*, genesis of RBC in health and progression of pathology in diseases^56^).

## Methods

### Protein purification

To dislodge different band 3-associated complexes from the human red blood cell membrane-cytoskeleton network, ghost membrane was sequentially treated with low-salt and high-salt buffers as reported before^8,57^ (Extended Data Fig. 1a). 50 mL packed human red blood cells (BioIVT) at 4°C were washed with five volumes of phosphate-buffered saline (PBS) at 2,000g for 10 min. All the following steps were performed at 4°C unless otherwise specified. The cells were then lysed in 10 volumes of hypotonic buffer containing 7.5 mM sodium phosphate at pH 7.5, 1 mM EDTA and protease inhibitors [0.5 mM phenylmethylsulphonyl fluoride (PMSF), 0.7 μg/mL pepstatin A, 2.5 μg/mL aprotinin, 5 μg/mL leupeptin] for 30 min. The lysate was centrifuged for 30 min at 20,000g to pellet the ghost membrane. The ghost membrane was further washed in the hypotonic buffer and pelleted at 20,000g for 30 min four times, followed by extraction in a low-salt buffer containing 0.1 M KCl, 7.5 mM sodium phosphate at pH 7.5, 1 mM EDTA, 1 mM dithiothreitol (DTT), 1% n-Dodecyl-beta-Maltoside (DDM) and protease inhibitors for 1 h. Subsequently, the sample was centrifuged at 20,000g for 20 min, resulting in the supernatant (low-salt fraction) and the pellet. The pellet was further extracted with a high-salt buffer containing 1 M KCl, 7.5 mM sodium phosphate at pH 7.5, 1 mM EDTA, 1 mM DTT, 1% DDM and protease inhibitors for 1 h. After centrifugation at 45,000g for 30 min, the supernatant (high-salt fraction) was obtained for further purification.

The low-salt and high-salt fractions were further purified by using gel-filtration column Superose 6 Increase 10/300 GL (GE Healthcare). The low-salt fraction was injected into the column in SEC150 buffer (10 mM Tris 7.5, 150 mM NaCl, 1 mM DTT, 0.015% DDM and protease inhibitors). After analysis by SDS-PAGE, band 3 fractions were pooled and purified by using the same column for a second time (Extended Data Fig. 1b). The peak fraction was concentrated and used for cryo-EM grid preparation. The high-salt fraction was injected into the gel-filtration column in SEC500 buffer (10 mM Tris 7.5, 500 mM NaCl, 1 mM DTT, 0.015% DDM and protease inhibitor), and resulted in an elution volume of 14.5 mL of high-salt fraction 1 (band 3-protein 4.2 complex) and 12.5 mL of high-salt fraction 2 (ankyrin complex) (Extended Data Fig. 1c). Band 3-protein 4.2 complex was pooled and further purified by using the same column for a second time in SEC300 buffer (10 mM Tris 7.5, 300 mM NaCl, 1 mM DTT, 0.015% DDM and protease inhibitor) (Extended Data Fig. 1d). The good fractions were combined and concentrated for cryo-EM. For band 3-protein 4.2 complex used in analytical gel filtration, pooled fractions from the first gel-filtration purification were further purified by anion exchange column (Source-15Q, GE Healthcare) and then subjected to a second gel-filtration column in SEC150 buffer. The ankyrin complex was stabilized by a Grafix^18,58^ method after the first gelfiltration purification of the high-salt fraction. Glycerol gradient was made by mixing 6 mL light buffer (20 mM HEPES 7.5, 300 mM NaCl, 0.015% DDM, 10% glycerol) and 6 mL heavy buffer [20 mM HEPES 7.5, 300 mM NaCl, 0.015% DDM, 30% glycerol, 0.2% glutaraldehyde (Polysciences)] in a gradient master (BioComp). Ankyrin complex was concentrated to 200 μL and dialyzed to buffer containing 20 mM HEPES 7.5, 300 mM NaCl, 0.015% DDM. The sample was loaded on top of the glycerol gradient and centrifuged at 4°C for 18 h at a speed of 35,000 rpm in SW-41Ti rotor (Beckman). Fractions of 500 μL were collected, and the cross-link reaction was quenched by adding Tris 7.5 to a final concentration of 50 mM. Good fractions after negative stain screening were combined and subjected to the gel-filtration column in SEC300 buffer (Extended Data Fig. 1e). Finally, the fractions in the 12.5 mL peak were collected and used for cryo-EM.

All recombinant proteins and mutants were overexpressed in *E. coli* strain BL21(DE3). DNA sequences encoding ankyrin ARs 1-24 (*a.a.* 1-827), ARs 13-24 (*a.a.* 402-827), ARs 16-24 (*a.a.* 494-827), ARs 17-24 (*a.a.* 527-827), ARs 18-24 (*a.a.* 561-827), ARs 19-24 (*a.a.* 597-827), ARs 20-24 (*a.a.* 630-827) and band 3 cytoplasmic domain (*a.a.* 1-379) were cloned into a modified pET-28a vector with an N-terminal hexahistidine tag followed by a SUMO tag. Mutants were generated by QuikChange mutagenesis and confirmed by DNA sequencing. Proteins were expressed in *E. coli* and induced with 0.5 mM isopropyl-β-D-thiogalactoside (IPTG, Sigma) at an OD_600_ of 0.8. The culture was incubated at 25°C overnight. Cells were harvested by centrifugation and resuspended in a buffer containing 20 mM Tris 7.5, 300 mM NaCl, 1 mM PMSF and 1 mM benzamidine. The suspensions were lysed by using a cell disruptor (Avestin). After high-speed centrifugation at 30,000g for 1 h, the supernatant was loaded to a column with HisPur cobalt resin (Thermo Fisher). After a wash step, proteins were eluted with a buffer containing 20 mM Tris 7.5, 150 mM NaCl,1 mM benzamidine and 250 mM imidazole. For band 3 cytoplasmic domain, the His6-SUMO tag was removed by ULP1 (a SUMO protease). For all the ankyrin constructs, the His6-SUMO tag was retained. Proteins were further purified by ion-exchange column (Source-15Q, GE healthcare) and polished by gel-filtration column (Superdex-200, GE Healthcare) in SEC150 buffer without protease inhibitors. The purified proteins were concentrated and stored at −80°C.

### Cryo-EM sample preparation and image acquisition

For cryo-EM sample optimization, an aliquot of 3 μL of sample was applied onto a glow-discharged holey carbon-coated copper grid (300 mesh, QUANTIFOIL® R 2/1) or holey gold grid (300 mesh, UltrAuFoils® R 1.2/1.3). The grid was blotted with Grade 595 filter paper (Ted Pella) and flash-frozen in liquid ethane with an FEI Mark IV Vitrobot. An FEI TF20 cryo-EM instrument was used to screen grids. Cryo-EM grids with optimal particle distribution and ice thickness were obtained by varying the gas source (air using PELCO easiGlow™, target vacuum of 0.37 mbar, target current of 15 mA; or H_2_/O_2_ using Gatan Model 950 advanced plasma system, target vacuum of 70 mTorr, target power of 50 W) and time for glow discharge, the volume of applied samples, chamber temperature and humidity, blotting time and force, and drain time after blotting. Our best grids for low salt fraction were obtained with holey carbon-coated copper grids, 20 s glow discharge using H_2_/O_2_ and with the Vitrobot sample chamber temperature set at 8°C, 100% humidity, 6 s blotting time, 3 blotting force, and 0 s drain time. The best grids for high-salt fraction 1 and 2 were obtained with holey gold grids, 20 s glow discharge using H_2_/O_2_ and with the Vitrobot sample chamber set at 8°C temperature, 100% humidity, 6 s blotting time, 3 blotting force, and 0 s drain time.

Optimized cryo-EM grids were loaded into an FEI Titan Krios electron microscope with a Gatan Imaging Filter (GIF) Quantum LS device and a post-GIF K2 or K3 Summit direct electron detector. The microscope was operated at 300 kV with the GIF energy-filtering slit width set at 20 eV. Movies were acquired using SerialEM^59^ by electron counting in super-resolution mode at a pixel size of 0.535 Å/pixel or 0.55 Å/pixel with a total dosage ~50 e^-^/Å^2^/movie. Image conditions are summarized in Table 1.

### Cryo-EM reconstruction

Frames in each movie were aligned for drift correction with the GPU-accelerated program MotionCor2 (ref.^60^). Two averaged micrographs, one with dose weighting and the other without, were generated for each movie after drift correction. The averaged micrographs have a calibrated pixel size of 1.062 Å (low-salt fraction) or 1.1 Å (high-salt fraction 1 and 2) at the specimen scale. The averaged micrographs without dose weighting were used only for defocus determination, and the averaged micrographs with dose weighting were used for all other steps of image processing. Workflows are summarized in Extended Data Fig. 2, 3 and 4 for low-salt fraction, high-salt fraction 1 and high-salt fraction 2, respectively.

For low-salt fraction (band 3), a total of 9,455 averaged micrographs were obtained, of which 1,000 micrographs were subjected to a quick analysis in cryoSPARC v3 (ref.^61^). The defocus values of the 9,455 averaged micrographs were determined by CTFFIND4 (ref.^62^). 2,658,315 particles were automatically picked without reference using Gautomatch (https://www2.mrc-lmb.cam.ac.uk/research/locally-developed-software/zhang-software/). Several rounds of reference-free 2D classification were subsequently performed in RELION3.1^63,64^ to remove “bad” particles (i.e., classes with fuzzy or un-interpretable features), yielding 961,892 good particles. To retrieve more real particles from the micrographs, Topaz^65^, a convolutional neural network-based particle picking software, was trained by the final coordinates from cryoSPARC and used for the second round of particle picking. 2,953,637 particles were obtained initially and resulted in 976,060 good particles after rounds of 2D classification in RELION. After the two sets of particles were combined and duplicates removed, a total of 1,286,729 particles were collected. A global search 3D classification in RELION was performed, with the map from cryoSPARC as the initial model. One good class containing 530,179 particles was selected and subjected to a final step of 3D auto-refinement in RELION. Membrane part and cytoplasmic part of band 3 are refined together, without local refinement. The two half-maps from this auto-refinement step were subjected to RELION’s standard post-processing procedure, yielding a final map with an average resolution of 4.8 Å.

For high-salt fraction 1 (band 3-protein 4.2 complex), a total of 20,842 averaged micrographs was obtained and subjected to particle picking in Gautomatch. 2,031,749 particles were selected after 2D and 3D classification in RELION. The coordinates of the selected particles were used for Topaz training in the second round of particle picking. 2,879,888 particles were selected after the second round of particle sorting. The two sets of selected particles were combined and duplicates removed, resulting in 3,864,165 particles. These particles were subjected to a global search 3D classification with K=5, resulting in four different structures of band 3-protein 4.2 complex: 1,048,826 particles (27.1%) in the class of band 3 with a loosely bound protein 4.2 (B_2_P_1_^loose^), 406,951 particles (10.5%) in the class of band 3 with protein 4.2 binding vertically (B_2_P_1_^vertical^), 246,059 particles (6.4%) in the class of band 3 with two protein 4.2 binding vertically (B_2_P_2_^vertical^) and 2,162,329 particles (56%) in the major class of band 3 with protein 4.2 binding diagonally (B_2_P_1_^diagonal^). The B_2_P_1_^loose^ complex, B_2_P_1_^vertical^ complex and B_2_P_2_^vertical^ complex were further 3D classified with the skip-align option in RELION and reconstructed to 4.1 Å, 4.6 Å and 4.6 Å, respectively. For the B_2_P_1_^diagonal^ complex, the structure was classified and refined to 3.6 Å with an overall mask. When masks for the cytoplasmic part and the membrane part were applied, the structures of the cytoplasmic part and the membrane part were reconstructed to 3.1 Å and 3.3 Å, respectively.

For high-salt fraction 2 (ankyrin complex), 21,187 good micrographs were obtained. Using a strategy similar to that of the band 3 dataset, a total of 4,048,034 unique particles were collected after two rounds of particle picking and 2D classification. The selected particles were subjected to 2D and 3D classification, and one good class containing 593,880 particles was selected. To further increase the number of good particles, a method of seed-facilitated 3D classification^66^ was used. Briefly, all the raw particles from autopick were divided into six subsets and then mixed with the good particles (seed) from the previous step of 3D classification. Subsequently, after applying 3D classification separately, all the good classes were collected and combined, followed by 2D and 3D classification to generate a total of 1,410,445 good particles. These particles were further 3D classified into four classes, resulting in two structures, with 383,995 particles in B_2_P_1_A_1_ complex (one ankyrin molecule binds to one B_2_P_1_^diagonal^ complex via protein 4.2) and 63,084 particles in B_2_P_1_A_2_ complex (two ankyrin molecules bind to one B_2_P_1_^diagonal^ complex via protein 4.2 and band 3, respectively). The two complexes were classified and refined in cryoSPARC using non-uniform refinement to resolutions of 4.4 Å and 5.7 Å for B_2_P_1_A_1_ complex (160,406 particles) and B_2_P_1_A_2_ complex (34,612 particles), respectively. A further step of focused refinement improved the core region’s resolutions to 4.1 Å and 4.6 Å for B_2_P_1_A_1_ complex and B_2_P_1_A_2_ complex, respectively. When lowering the threshold of the maps, smeared density emerges at the edges of the box, indicating that the box size of 384 pixels is not big enough to include all the densities. Therefore, these particles were re-centered and extracted in a box size of 560 pixels. After 3D classification and refinement, two structures of ankyrin-containing complex were reconstructed to 5.6 Å and 8.5 Å for B_4_P_1_A_1_ complex and (B_2_P_1_A_1_)_2_ complex, respectively.

### Resolution assessment

All resolutions reported above are based on the “gold-standard” FSC 0.143 criterion^67^. FSC curves were calculated using soft spherical masks, and high-resolution noise substitution was used to correct for convolution effects of the masks on the FSC curves^67^. Prior to visualization, all maps were sharpened by applying a negative B factor, estimated using automated procedures^67^. Local resolution was estimated using ResMap^68^. The overall quality of the maps for band 3, band 3-protein 4.2 complexes, and ankyrin-containing complexes is presented in Extended Data Fig. 2d-f, 3d-h, and 4d-f, respectively. The reconstruction statistics are summarized in Table 1.

### Atomic modeling, model refinement and graphics visualization

Atomic model building started from the B_2_P_1_^diagonal^ complex map, which had the best resolution. We took advantage of the reported crystal structure of mdb3 (PDB: 4YZF)^16^, which was fitted into the focus-refined membrane domain map (3.3 Å) by UCSF Chimera^69^. We manually adjusted its side chain conformation and, when necessary, moved the main chains to match the density map using Coot^70^. This allowed us to identify extra densities for loop N554-P566, loop A641-W648 and N-acetylglucosamine (NAG) close to residue N642, as well as lipids at the dimer interface that were tentatively assigned as DDM or cholesterol accordingly. For the cytoplasmic part, the crystal structure of cdb3 (PDB: 1HYN)^15^ was fitted into the focus-refined cytoplasmic domain map (3.2 Å) and manually adjusted. This enabled us to identify the extra densities for the N-terminus of the cdb3 (*a.a.* 30-55), absent in the crystal structure and interacting with protein 4.2 in our B_2_P_1_^diagonal^ complex. Next, we built the atomic model for protein 4.2 *de novo.* Protein sequence assignment was mainly guided by visible densities of amino acid residues with bulky side chains, such as Trp, Tyr, Phe, and Arg. Other residues including Gly and Pro also helped the assignment process. Unique patterns of sequence segments containing such residues were utilized for validation of residue assignment. Finally, the models of the membrane part and the cytoplasmic part were docked into the 3.6 Å overall map in Chimera. As the map resolution of the CM-linker between cdb3 and mdb3 is insufficient for *de novo* atomic modeling, we traced the main chain using Coot for *a.a.* 350-369.

For the structure of the band 3 dimer, models of cdb3 and mdb3 from the B_2_P_1_^diagonal^ complex were fitted into the cryo-EM map in Chimera and manually adjusted using Coot. For the structures of B_2_P_1_^vertical^ and B_2_P_2_ ^vertical^, models of mdb3, cdb3 and protein 4.2 from the B_2_P_1_^diagonal^ complex were docked and manually adjusted.

For the structure of the B_2_P_1_A_1_ complex, we first docked the model of the B_2_P_1_^diagonal^ complex into the cryo-EM map. Ankyrin repeats 6-20 (*aa.* 174-658) with side chains were built *de novo* using the focus map at 4.1 Å. ARs 21-24 were assigned with guidance from previous crystal structures of ankyrins (ARs 1-24 of AnkyrinB, PDB: 4RLV^40^; ARs 13-24 of AnkyrinR, PDB:1N11^17^) and truncated to Cβ due to the lack of side-chain densities. Next, the structure of B_2_P_1_A_1_ complex was fitted into the maps of B_2_P_1_A_2_. This enabled us to identify extra densities for a second ankyrin which directly interacts with band 3. The bulky side chains of the second ankyrin were built using the focus map at 4.6 Å. The assignment of the band 3-associated ankyrin was further verified in both the B_4_P_1_A_1_ and (B_2_P_1_A_1_)_2_ complexes.

The atomic models were refined using PHENIX^71^ in real space with secondary structure and geometry restraints. All the models were also evaluated using the wwPDB validation server (Table 1). Representative densities are shown in Extended Data Fig. 2g, 3i and 4g-h. Visualization of the atomic models, including figures and movies, was accomplished in UCSF Chimera and Chimera X^69^.

### Structure-guided mutagenesis and pull-down assay

The pull-down assay was performed with HisPur cobalt resin at 4°C in a binding buffer containing 10 mM Tris 7.5, 150 mM NaCl, 0.015% DDM. 20 μL cobalt resin was used in a 200 μL binding reaction. Recombinant proteins were used in this assay. His6-SUMO tagged ankyrin at a final concentration of 4 μM was pre-incubated with the resin and then mixed with 8 μM of band 3 cytoplasmic domain or mutant. After 60 min of incubation, the resin was washed four times with 1 mL of the binding buffer containing 10 mM imidazole and eluted with the binding buffer containing 250 mM imidazole. Samples were analyzed by SDS-PAGE and stained with InstantBlue (abcam). All experiments were repeated at least three times.

### Analytical gel filtration

Analytical gel filtration chromatography was carried out with the Superose 6 Increase 10/300 GL column at 4°C. The column was equilibrated with SEC150 buffer without protease inhibitors. Band 3-protein 4.2 complex was purified from the erythrocyte membrane. Ankyrin and band 3 cytoplasmic domain were purified from *E. coli.* The His6-SUMO tag of ankyrin was retained. A protein sample of 500 μL was injected into the column and eluted at a flow rate of 0.5 mL/min. Fractions were then analyzed by SDS-PAGE.

### Reporting summary

Further information on research design is available in the Nature Research Reporting Summary linked to this article.

## Data availability

Cryo-EM density maps have been deposited in the Electron Microscopy Data Bank under accession numbers EMD-26148 (band 3 dimer), EMD-26145 (B_2_P_1_^loose^), EMD-26146 (B_2_P_1_^vertical^), EMD-26147 (B_2_P_2_^vertical^), EMD-26142 (B_2_P_1_^diagonal^), EMD-26143 (membrane part of B_2_P_1_^diagonal^), EMD-26144 (cytoplasmic part of B_2_P_1_^diagonal^), EMD-26149 (B_2_P_1_A_1_), EMD-26150 (cytoplasmic part of B_2_P_1_A_1_), EMD-26151 (B_2_P_1_A_2_), EMD-26152 (focused refinement of B_2_P_1_A_2_), EMD-26153 (B_4_P_1_A_1_) and EMD-26154 [(B_2_P_1_A_1_)_2_]. Model coordinates have been deposited in the Protein Data Bank under accession numbers 7TW2 (band 3 dimer), 7TW0 (B_2_P_1_^vertical^), 7TW1 (B_2_P_2_^vertical^), 7TVZ (B_2_P_1_^diagonal^), 7TW3 (B_2_P_1_A_1_), 7TW5 (B_2_P_1_A_2_) and 7TW6 (B_4_P_1_A_1_). All other data needed to evaluate the conclusions in the paper are present in the paper and/or the supplementary materials.

## Acknowledgments

We thank Titania Nguyen, James Zhen and Alex Stevens for editorial assistance. This project is supported by grants from the US NIH (R01GM071940 to Z.H.Z.). We acknowledge the use of resources at the Electron Imaging Center for Nanomachines supported by UCLA and grants from the NIH (1S10OD018111 and 1U24GM116792) and the National Science Foundation (DBI-1338135 and DMR-1548924).

## Author contributions

Z.H.Z. conceived the project. X.X. and S.L. prepared samples, acquired and analyzed cryo-EM data. X.X. engineered and isolated the recombinant proteins, and performed biochemistry analyses. S.L. and X.X. built the models. X.X., S.L. and Z.H.Z. interpreted the results and wrote the manuscript.

---

**Extended Data Fig. 1:**
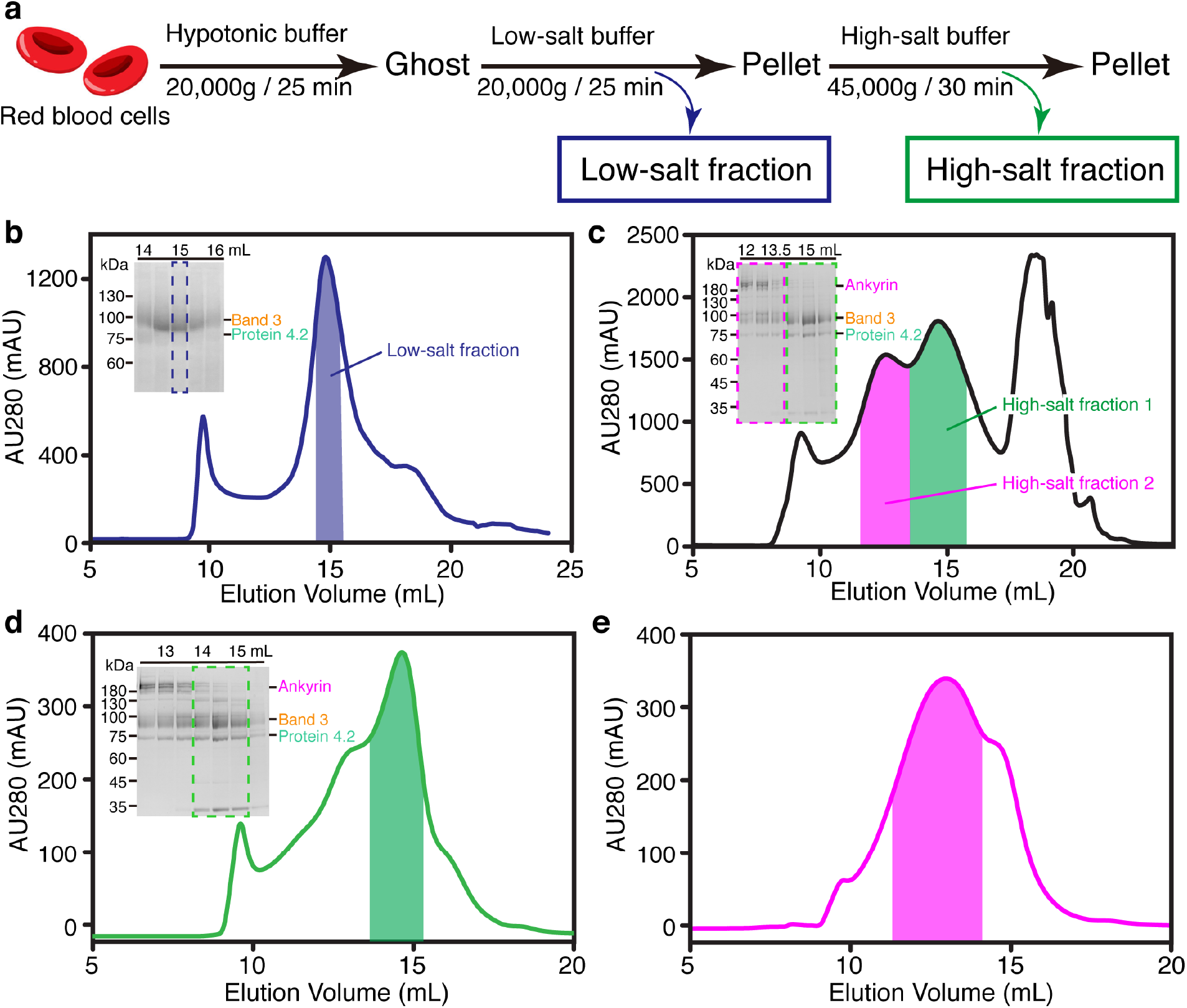
Purification and cryo-EM reconstruction of the erythrocyte membrane proteins. (**a**) Workflow of the stepwise fractionation of erythrocyte membrane proteins. (**b**) The second gel-filtration chromatography profile of the low-salt fraction. The result from SDS-PAGE analysis of the peak fractions is inserted in the upper left corner. The peak fractions were applied to SDS-PAGE and visualized by Coomassie blue staining. Dashed blue box on the gel and blue bar on the chromatogram indicate the fractions collected for cryo-EM. (**c**) The first gel-filtration chromatography profile of the high-salt fraction. Green and magenta boxes on the gel and green and magenta bars on the chromatogram indicate the fractions collected for the protein 4.2 complex and ankyrin complex, respectively. (**d**) The second gel-filtration chromatography profile and corresponding gel of the protein 4.2 complex. Dashed green box on the gel and green bar on the chromatogram indicate the fractions collected for cryo-EM. (**e**) The gel-filtration chromatography profile of the ankyrin complex after Grafix purification. Magenta bar on the chromatogram indicates the fractions collected for cryo-EM.

**Extended Data Fig. 2:**
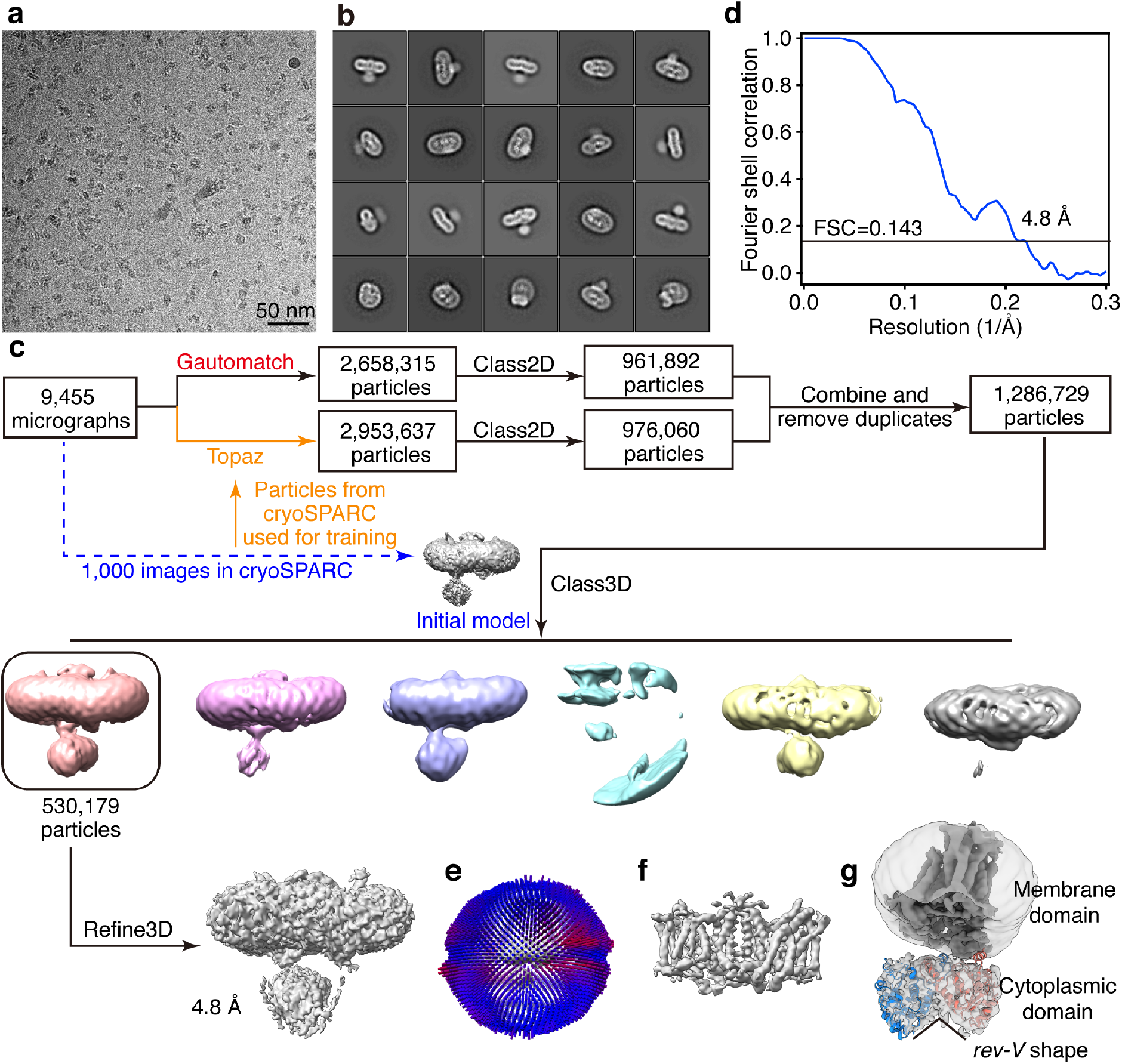
Cryo-EM analysis of the low-salt fraction (band 3). (**a**) Representative cryo-EM image of the low-salt fraction. (**b**) Selected 2D class averages of the cryo-EM particle images. (**c**) Flow chart of cryo-EM data processing. (**d**) Gold-standard Fourier shell correlation (FSC) curve for 3D reconstruction. (**e**) Angular distribution of cryo-EM reconstructions used for final refinement. (**f**) Density of the membrane domain. (**g**) Atomic model of band 3 cytoplasmic domain fitted into the cryo-EM density. A lower map threshold is used in (g) compared to that of (f) to better present the cytoplasmic domain.

**Extended Data Fig. 3:**
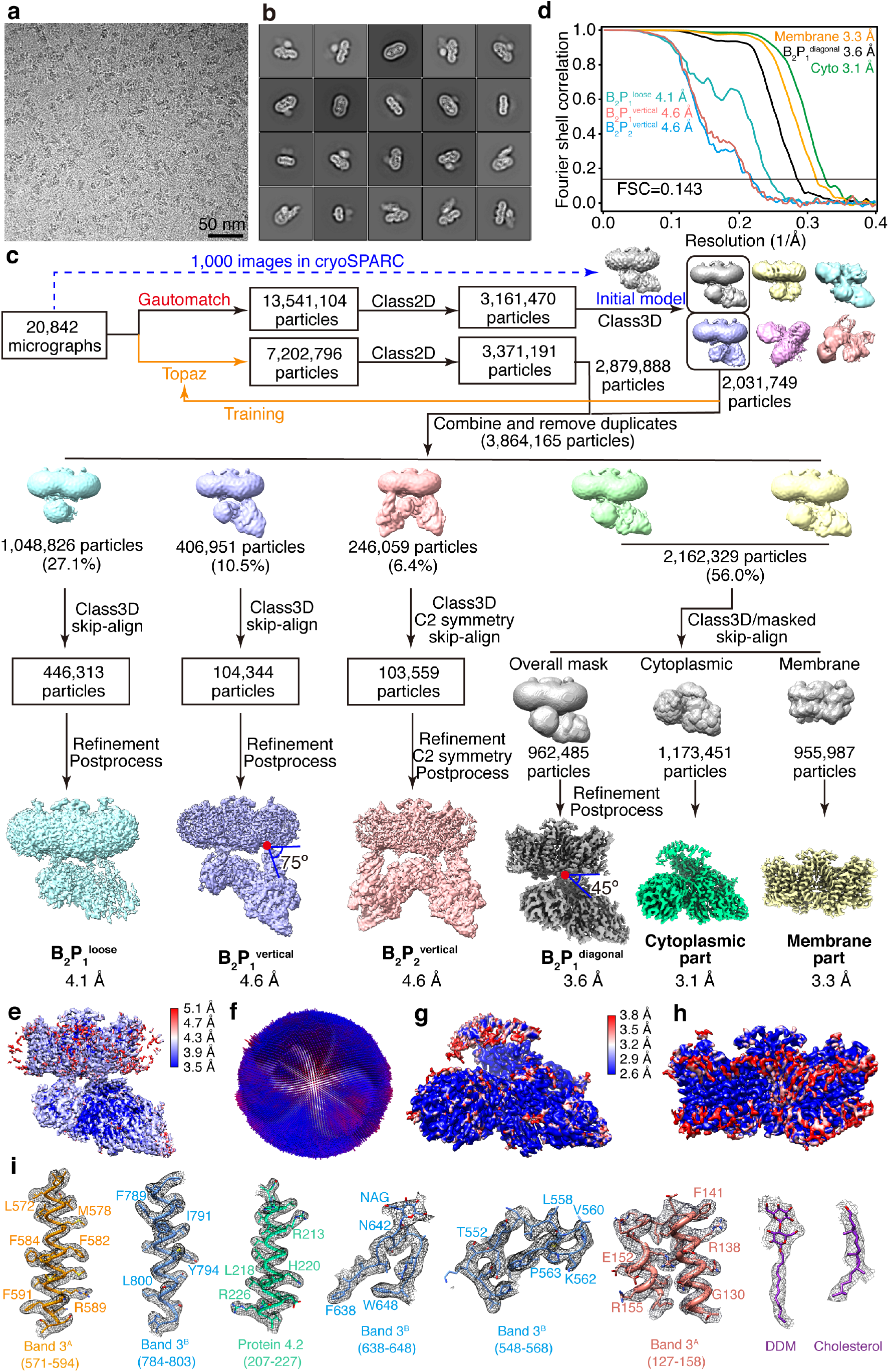
Image processing for the cryo-EM data of the high-salt fraction 1 (band 3-protein 4.2 complex). (**a**) Representative cryo-EM image of the high-salt fraction 1. (**b**) Selected 2D class averages of the cryo-EM particle images. (**c**) Flow chart of cryo-EM data processing. (**d**), Gold-standard Fourier shell correlation (FSC) curves for 3D reconstructions. (**e**) Local resolution of the overall map of B_2_P_1_diagonal complex. (**f**) Angular distribution of cryo-EM reconstruction of B_2_P_1_diagonal complex used for final refinement. (**g-h**) Local resolutions of the focus refinement maps of the cytoplasmic part and membrane part of B_2_P_1_diagonal complex. (**i**) Representative cryo-EM density maps of the B_2_P_1_diagonal complex.

**Extended Data Fig. 4:**
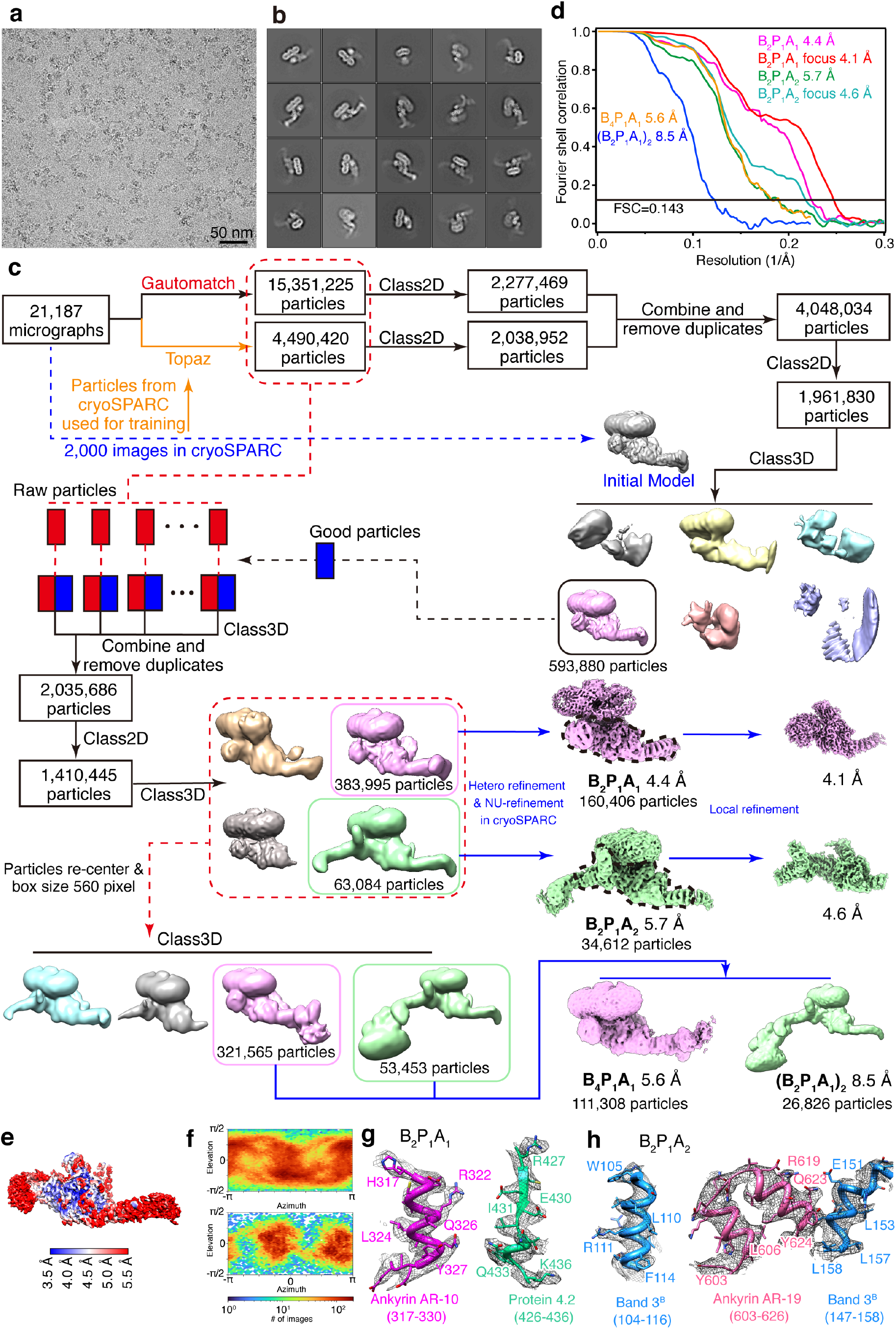
Cryo-EM analysis of the high-salt fraction 2 (ankyrin complex). (**a**) Representative cryo-EM image of the high-salt fraction 2. (**b**) Selected 2D class averages of cryo-EM particle images. (**c**) Flow chart of cryo-EM data processing. (**d**) Gold-standard Fourier shell correlation (FSC) curves for 3D reconstructions. (**e**) Local resolution of the overall map of B_2_P_1_A_1_ complex. **(f**) Angular distribution of cryo-EM reconstruction of B_2_P_1_A_1_ complex. (**g**) Representative cryo-EM density maps of the B_2_P_1_A_1_ complex showing the fragments of ankyrin and protein 4.2 at their binding interface. **(h**) Representative cryo-EM density maps of the B_2_P_1_A_2_ complex showing the fragments of ankyrin and band 3 at their binding interface.

**Extended Data Fig. 5:**
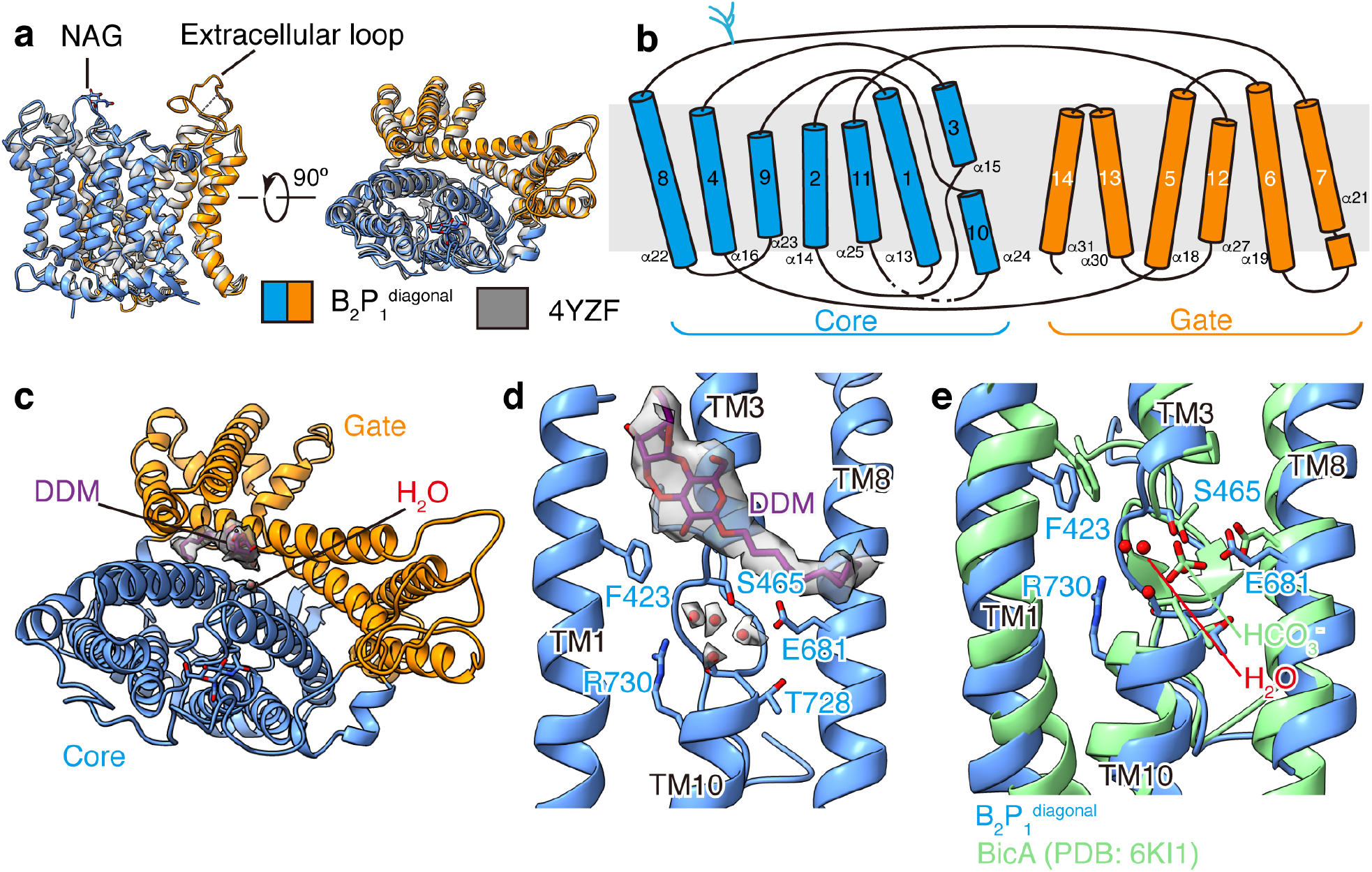
Structural analysis of the band 3 membrane domain. (**a**) Superposition of the band 3 membrane domain in B_2_P_1_^diagonal^ complex and reported crystal structure (PDB: 4YZF)^16^. (**b**) Topology of the transmembrane helices of band 3. (**c**) Density of the DDM molecule at the interface of the core and gate domain. (**d**) Enlarged view of the substrate binding site in B_2_P_1_^diagonal^ complex. Four water molecules were tentatively modelled into the cryo-EM density of band 3 near the substrate binding site. (**e**) Comparison of the substrate binding site in band 3 with that in bicarbonate transporter BicA (PDB: 6KI1)^25^.

**Extended Data Fig. 6:**
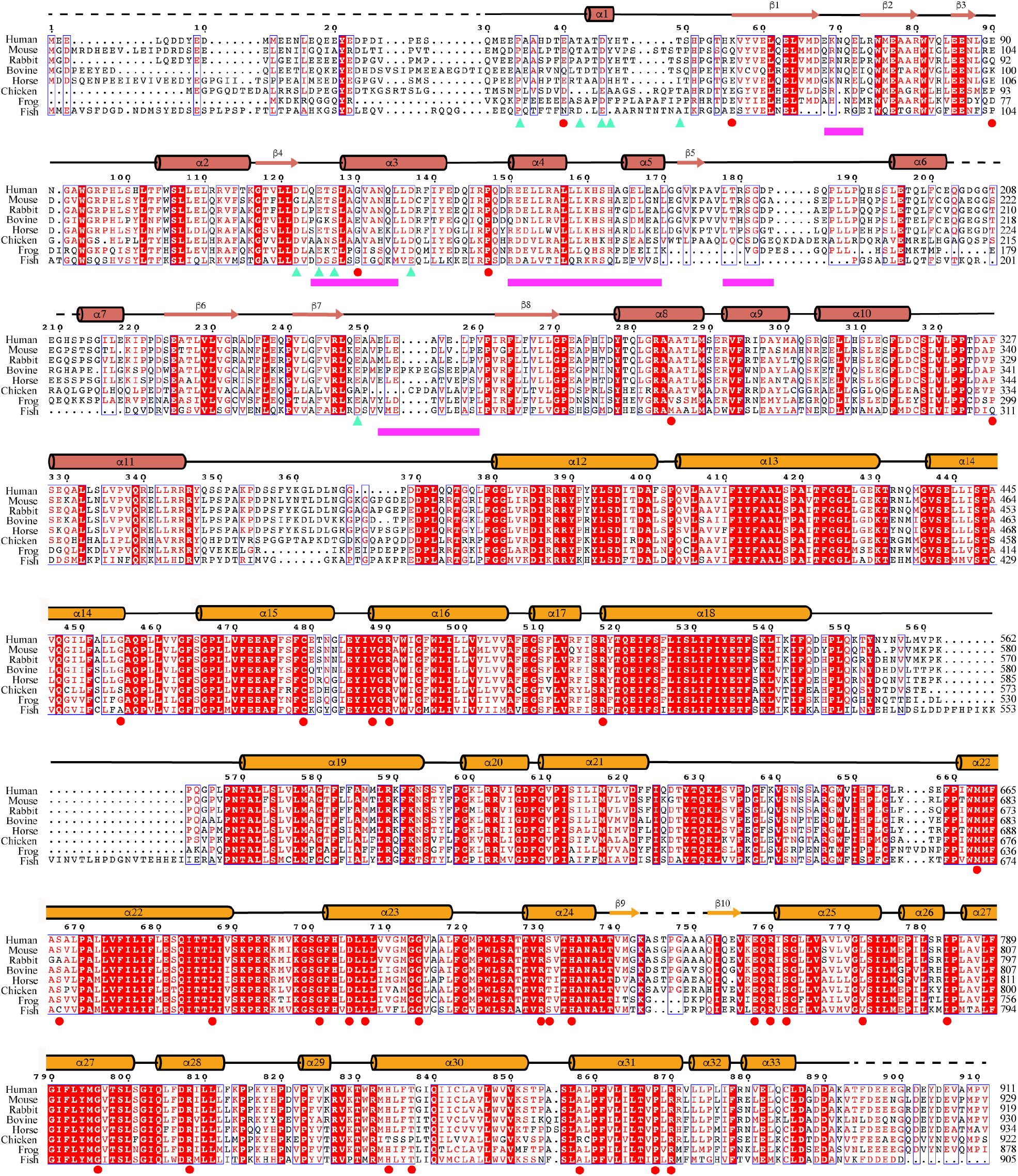
Sequence alignment of band 3 from different species. Sequences of human band 3 (P02730), mouse band 3 (P04919), rabbit band 3 (G1SLY0), bovine band 3 (Q9XSW5), horse band 3 (Q2Z1P9), chicken band 3 (P15575), frog band 3 (F6XSL8) and fish band 3 (Q7ZZJ7). The sequence alignment is done using the Clustal Omega server^72^; the figure is generated by ESPript 3^73^. Cyan triangles represent the band 3 residues interacting with protein 4.2; magenta bars indicate the regions of band 3 interacting with ankyrin; reported disease mutations on human band 3 are labeled as red circles.

**Extended Data Fig. 7:**
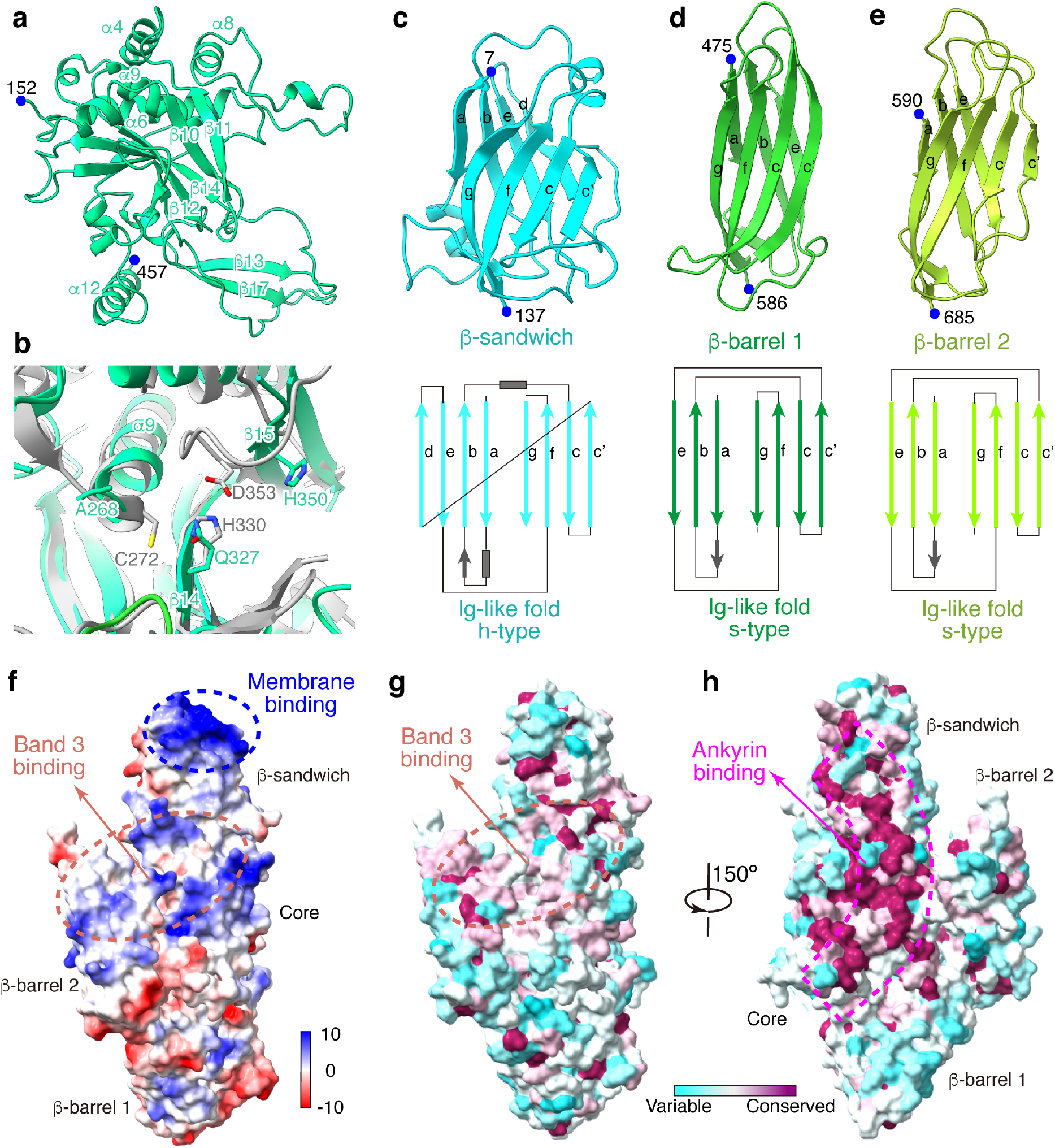
Structure of protein 4.2. (**a**) Structure of the core domain shown in ribbon. Residue numbers of its N and C terminus are labeled. **(b**) Superposition of protein 4.2 with transglutaminase (gray, PDB: 1L9N)^31^, showing the missing catalytic triad in protein 4.2. (**c-e**) Structures of the three Ig-like domains and illustrations of their secondary structure. (**f**) The electrostatic surface of protein 4.2, showing its membrane binding site (blue dashed circle) and band 3 binding interface (orange dashed circle). (**g-h**) Sequence conservation of protein 4.2 among mammals mapped to the structure. Orientation in (g) is the same as that in (f). Orange dashed circle shows the band 3 binding interface; magenta dashed box shows the ankyrin binding interface.

**Extended Data Fig. 8:**
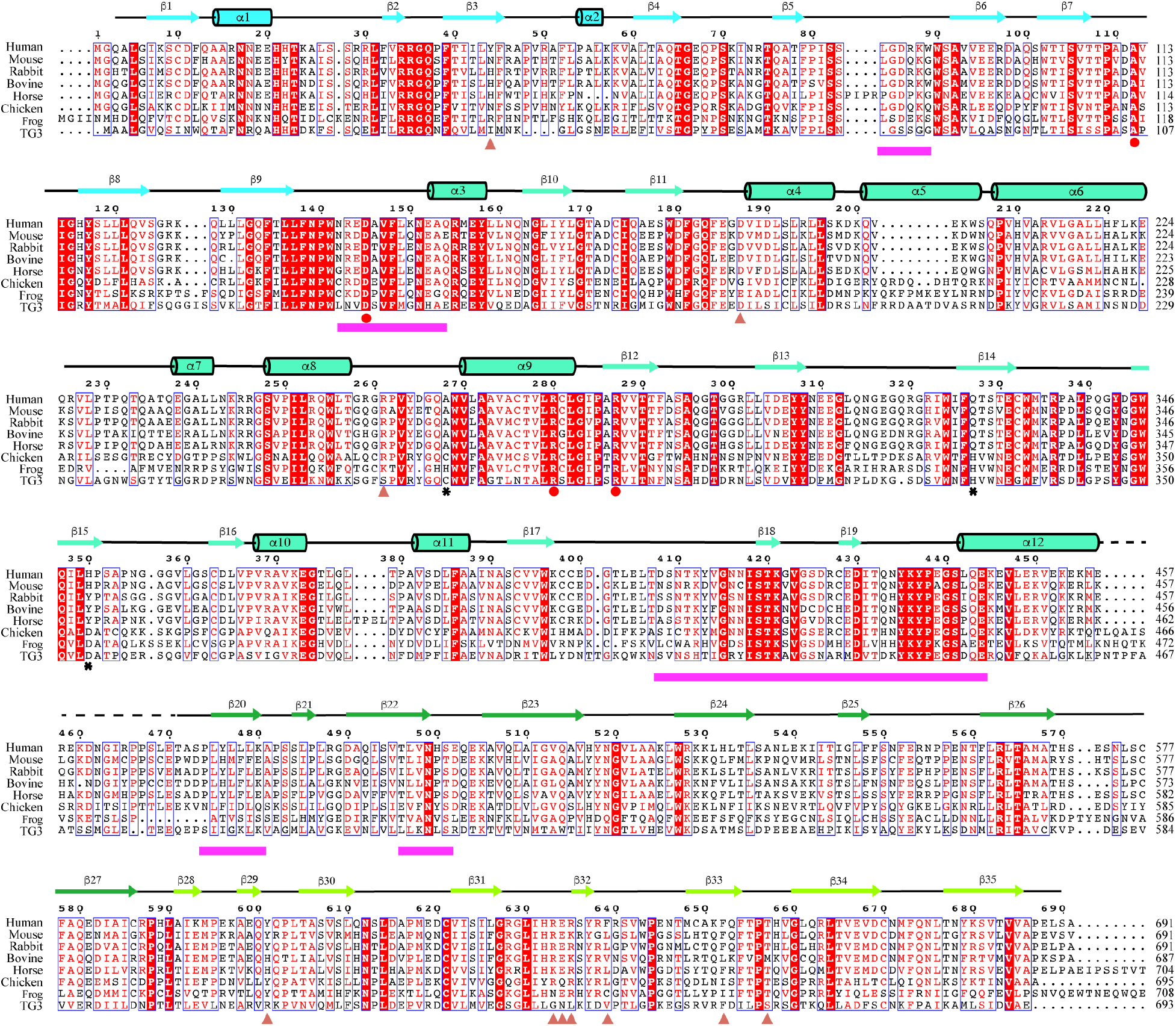
Sequence alignment of protein 4.2 from different species. Sequences of human protein 4.2 (P16452), mouse protein 4.2 (P49222), rabbit protein (G1TDR3), bovine protein 4.2 (O46510), horse protein 4.2 (F6ZDW1), chicken protein 4.2 (E1BQZ4), frog protein 4.2 (XP_018090678.1) and human transglutaminase 3 (Q08188). The sequence alignment is done using the Clustal Omega server^72^; the figure is generated in ESPript 3^73^. Salmon triangles represent protein 4.2 residues interacting with band 3; magenta bars indicate the regions of protein 4.2 interacting with ankyrin; reported disease mutations on human protein 4.2 are labeled as red circles; black stars indicate the catalytic residues of human transglutaminase 3.

**Extended Data Fig. 9:**
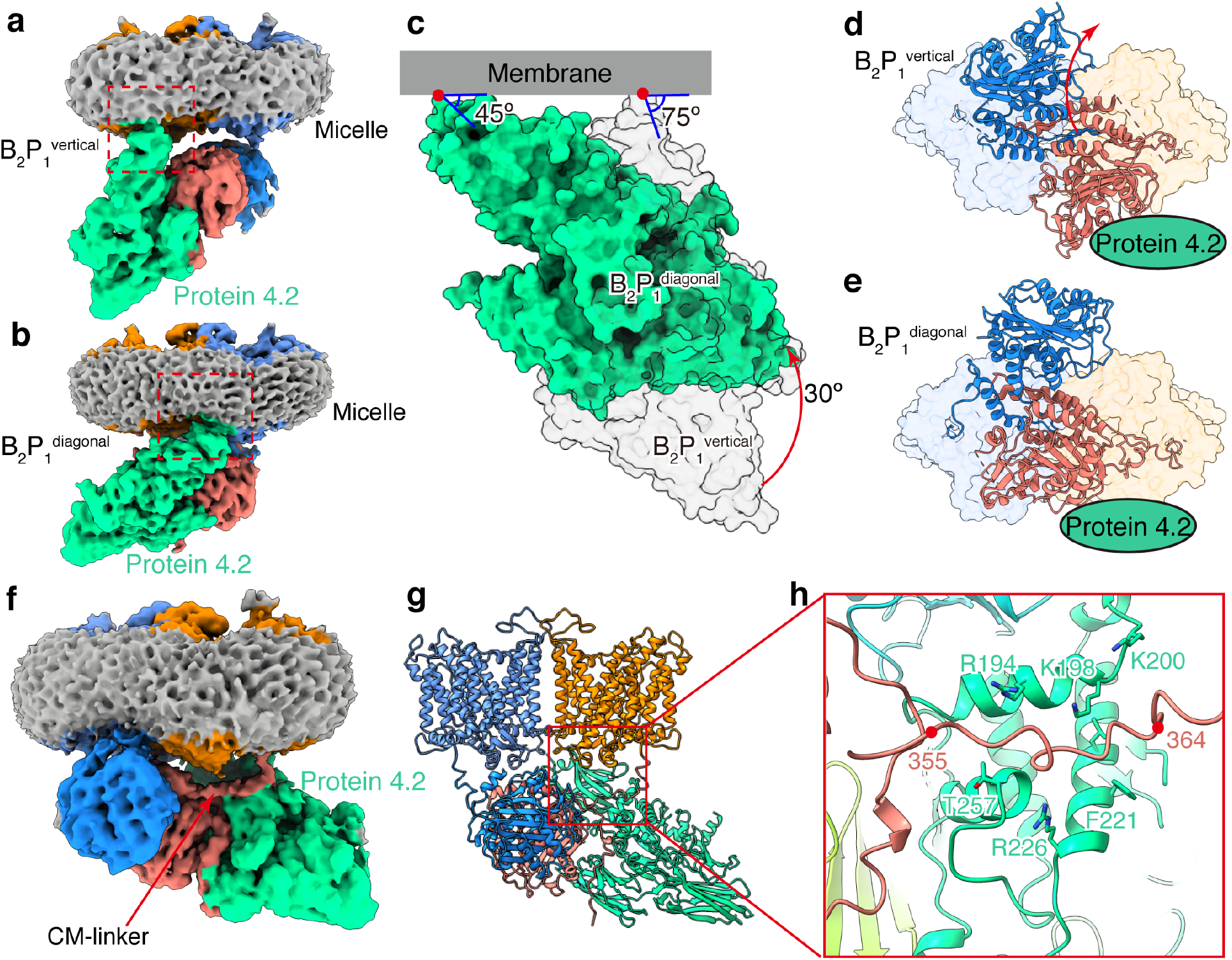
Conformational changes of band 3 and protein 4.2 during the assembly process. (**a-b**) Density map of the B_2_P_1_^vertical^ complex and B_2_P_1_^diagonal^ complex. Red box indicates the anchorage site of protein 4.2 N-termini to the membrane. (**c**) Rotation of protein 4.2 (red arrow) from vertical (transparent grey surface) to diagonal conformation (green surface). The two complexes are superposed according to the membrane domains of band 3. The cytoplasmic domains of band 3 are omitted for clarity. Angles between the membrane (grey bar) and protein 4.2 are labeled. (**d-e**) Rotation of the cytoplasmic domain of band 3 (red arrow) from B_2_P_1_^vertical^ to B_2_P_1_^diagonal^ complex viewed from the cytoplasmic side. The membrane domain of band 3 is shown as transparent surface and cytoplasmic domain as ribbon. Protein 4.2 is indicated as a green oval for clarity. (**f**) Density map of the B_2_P_1_^diagonal^ complex sharpened with B-factor of −50 Å^2^ showing the interaction of the CM-linker with protein 4.2. (**g-h**) Ribbon representation of the CM-linker region. Residues of protein 4.2 interacting with the CM-linker are labeled.

**Extended Data Fig. 10:**
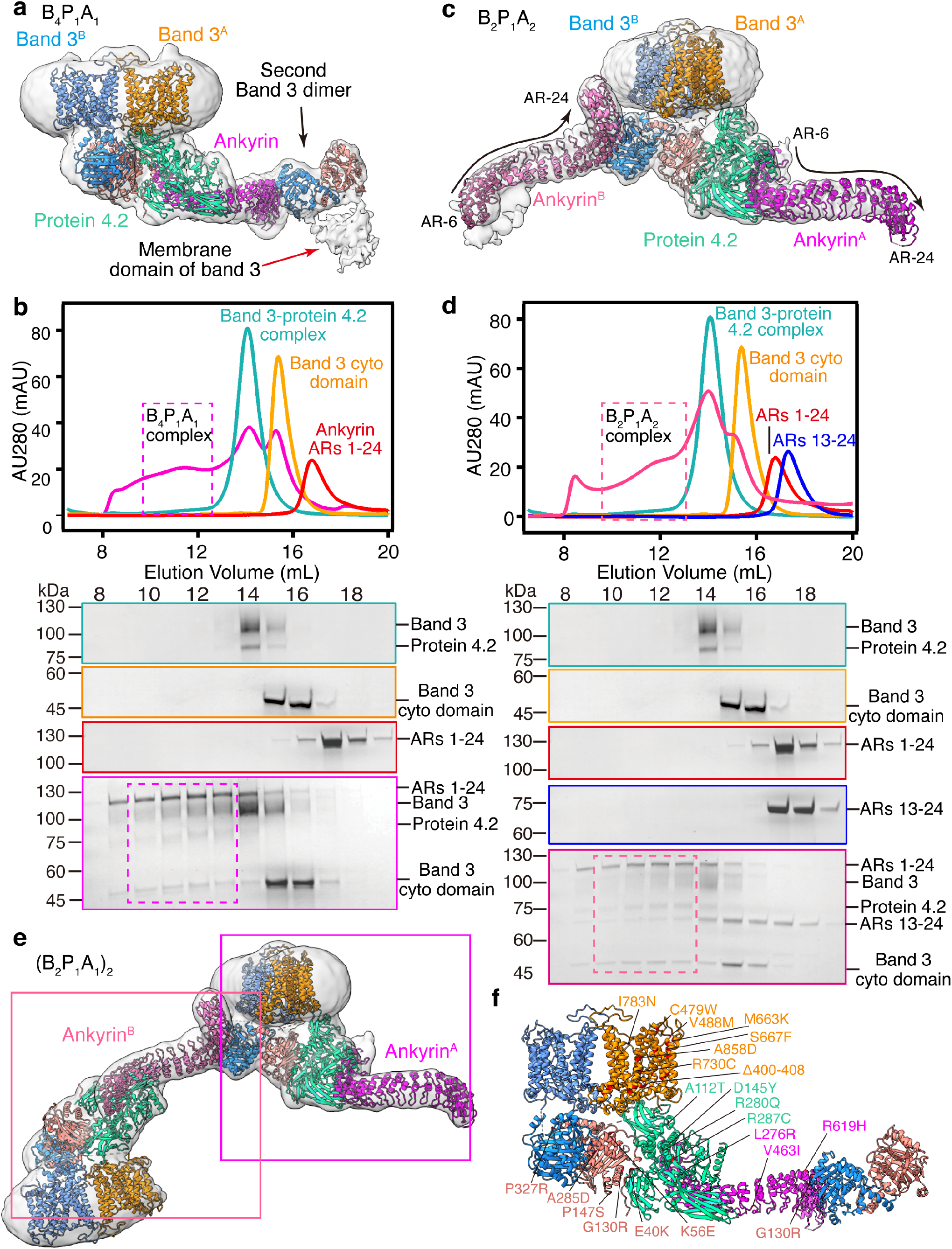
Analysis of the ankyrin-containing complexes. (**a**) Atomic model of B_4_P_1_A_1_ complex in ribbon superposed with the density map at a low threshold. Red arrow indicates the density of the membrane domain of the second band 3 dimer. (**b**) Analytical gel filtration assay showing the assembly of B_4_P_1_A_1_ complex in vitro. Dashed boxes show the position of B_4_P_1_A_1_ complex. Experiments were repeated for two times with similar results. (**c**) Atomic model of B_2_P_1_A_2_ complex in ribbon superposed with the density map at a low threshold. The density for the second band 3 dimer is indicated by arrow. (**d**) Analytical gel filtration assay showing the assembly of B_2_P_1_A_2_ complex in vitro. Dashed boxes show the position of B_2_P_1_A_2_ complex. The gel-filtration and SDS-PAGE results of protein 4.2 complex, band 3 cytoplasmic domain and ARs 1-24 are the same as that in (b). Second band 3 dimer may be incorporated into this complex, resulting in the B_4_P_1_A_2_. Experiments were repeated for two times with similar results. (**e**) Atomic model of (B_2_P_1_A_1_)_2_ complex in ribbon superposed with the density map. Boxes show the position of individual B_2_P_1_A_1_ complexes. (**f**) Reported disease mutations mapped on the structure of B_4_P_1_A_1_ complex.

**Movie S1.**
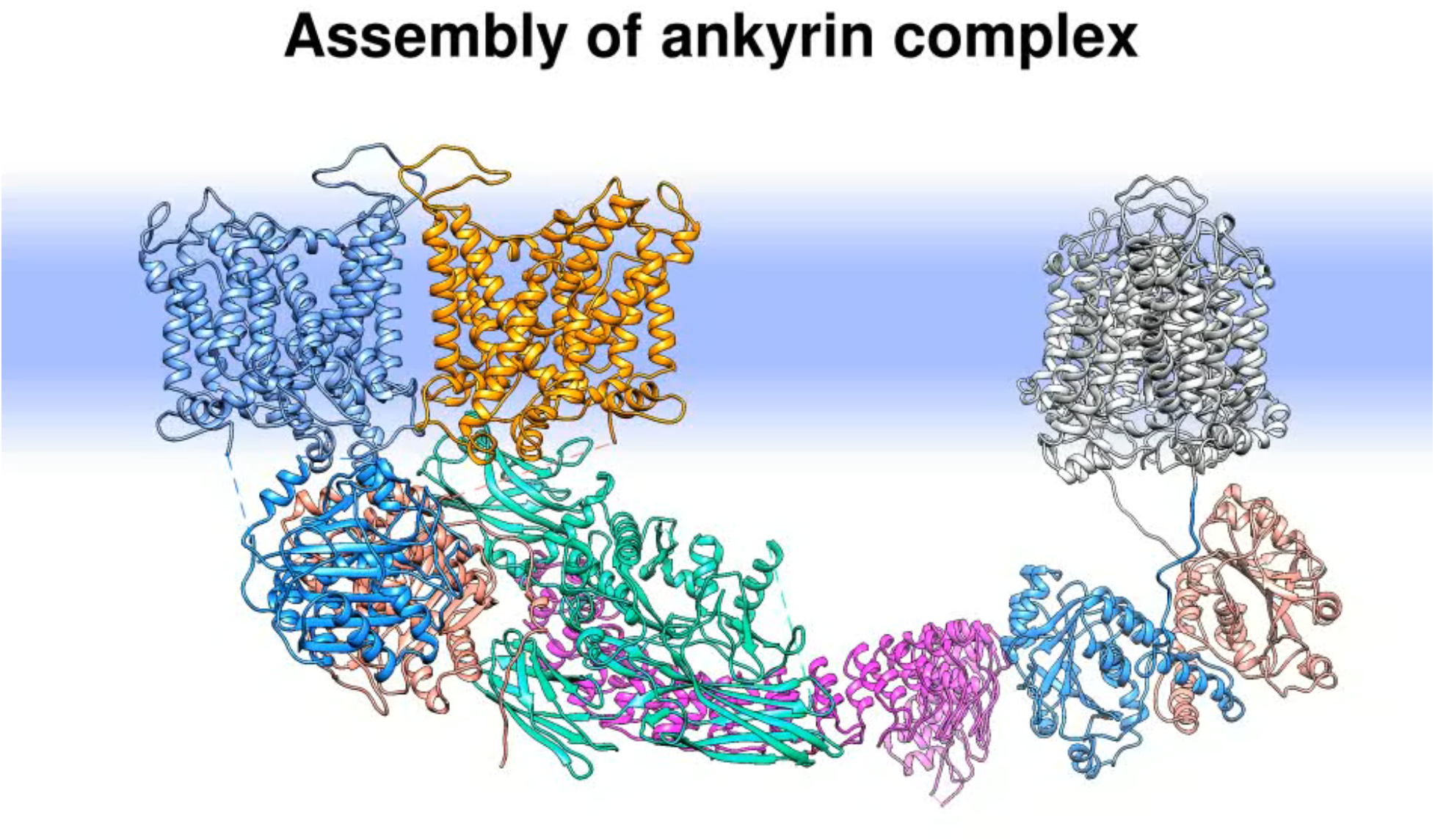
Sequential assembly of the ankyrin complex.

